# Long term AAV2/9-mediated silencing of PMP22 prevents CMT1A disease in rats and validates skin biomarkers as treatment outcome measure

**DOI:** 10.1101/2020.01.29.924605

**Authors:** Benoit Gautier, Helene Hajjar, Sylvia Soares, Jade Berthelot, Marie Deck, Scarlette Abbou, Graham Campbell, Claire-Maelle Fovet, Vlad Schütza, Antoine Jouvenel, Cyril Rivat, Michel Zerah, Virginie François Le Ravazet, Caroline Le Guiner, Patrick Aubourg, Robert Fledrich, Nicolas Tricaud

## Abstract

Charcot-Marie-Tooth disease 1A (CMT1A) results from a duplication of the *PMP22* gene leading to an excess of PMP22, a deficit of myelination and an instability of the myelin sheath in peripheral nerves. Patients present with reduced nerve conduction velocity, muscle waste, hand and foot deformations and foot drop walking problems. As gene silencing therapy has been shown to be effective in other monogenic neurological disorders, we evaluated the safety and efficacy of recombinant adeno-associated viral vector serotype 9 (AAV2/9)-based gene therapy for CMT1A. AAV2/9-mediated delivery of eGFP and shRNAs targeting PMP22 mRNA in the sciatic nerve allowed widespread gene expression in myelinating Schwann cells in mouse, rat and nonhuman primate. The treatment restored wild-type PMP22 level, increased myelination and prevented motor and sensory impairment over 12 months in a rat model of CMT1A. Intra-nerve injection limited off-target transduction and immune response to barely detectable levels. A combination of previously characterized human skin biomarkers successfully discriminated treated animals from their untreated littermate controls indicating their potential use as part of outcome measures in future clinical trials. Our results support intra nerve injection of AAV2/9 as an effective strategy for the treatment of CMT1A as well as other demyelinating CMT diseases.

## Introduction

Peripheral nerves that bundle axons emanating from neuronal cell bodies are found throughout the body forming the peripheral nervous system (PNS). These axons are covered in Schwann cells and those of large calibre are wrapped in a myelin sheath made by myelinating Schwann cells (mSC) to assure an extremely fast nerve conduction (up to 100 m/s). The myelin forms several successive segments named internodes, which electrically isolate the axonal membrane except at unmyelinated nodes of Ranvier where the action potentials are propagated [52].

This myelin sheath is critical for both motor and sensory functions in humans. Indeed, several peripheral nerve diseases impair the myelin sheath leading to reduced nerve conduction velocity, nerve dysfunction, muscle waste, limb extremities deformations and walking and sensory problems [6]. The large majority of patients suffering from hereditary diseases of peripheral nerves, namely Charcot-Marie-Tooth diseases (CMT), have defects in the myelin sheath formation, function or maintenance [10]. The most common of these myelin-related CMT diseases is CMT1A (prevalence 5-10/10000) [47]. This disease, resulting from the duplication of the *PMP22* gene, is characterized by a large heterogeneity of symptoms, the most serious being feet and hands deformation and walking problems [44]. PMP22 protein is a transmembrane glycoprotein located in the myelinated internode. In CMT1A, PMP22 protein overexpression in Schwann cells leads to defects in the number of myelinated segments that are developed, short internodes, myelin sheath defects, myelin degeneration (demyelination) and finally axonal loss [35].

No cure exists for this disease, however, pharmacological treatments have been investigated recently [44]. Some preclinical studies in rodents demonstrated that antisense oligonucleotides targeting PMP22 mRNA expression significantly improved the phenotype [33, 34, 66], suggesting PMP22 silencing is an effective way to tackle the disease. Intraperitoneal injection of human recombinant soluble Neuregulin-1 that promotes myelination in a rat model of CMT1A increased myelination, prevented axonal loss and dramatically delayed the occurrence of the disease [16, 37]. However, in most cases, the benefit only lasted as long as the treatment was administrated.

Gene therapy therefore constitutes an alternative therapy allowing for long term benefits for the patients [17]. Indeed, proofs of concept for gene therapy using lentiviral vectors injected in the spinal cord of myelin-related CMT mouse models have been reported [29, 30, 55]. Nevertheless, currently, *in vivo* gene therapy assays mostly use adeno-associated virus (AAV)-based strategies as these vectors do not integrate the genome of transduced cells, spread more easily in the tissues and display a limited immunogenicity [9]. AAV serotypes 2/9 and 2/rh10 are commonly used to transduce the central nervous system and several clinical trials are ongoing with promising results [17, 27]. Moreover, an AAV9-based therapy recently obtained market authorization from FDA to treat spinal motor atrophy in infants [38]. Gene therapy may therefore represent an efficient and safe way to treat CMT1A in the long term. However, the pattern of transduction of Schwann cells with AAV2/9 or rh10 remains unclear.

Recent clinical trials for peripheral neuropathies, and in particular for CMT1A, have shown that the chronicity of the disease makes the evaluation of the treatment outcome very difficult. Composite scores have been shown to be the most reliable outcome measure, but they remain poorly discriminant [36, 41, 49]. Molecular biomarkers have been characterized for CMT1A and some of them allow discrimination on the basis of disease severity [14]. However, none of them has been validated yet as outcome measure for a therapy.

Here we show that myelinating Schwann cells of mouse, rat and nonhuman primate sciatic nerves are widely and specifically transduced by AAV2/9 when injected directly into the nerve. Using an AAV2/9 expressing a small hairpin inhibitory RNA (shRNA) directed against PMP22 mRNA, we treated a rat model of CMT1A when myelination begins. A single injection treatment in both sciatic nerves prevented the disease symptoms for at least one year. The combined expressions of human biomarkers in the paw skin of rats correlated with the phenotype and allowed discrimination between treated animals and their sham-treated littermates, indicating that these markers can be used as an outcome measure of the treatment. In addition, the dispersion of the vector remained limited to the injected nerves and the humoral immune response generated against the vector remained barely detectable in injected animals. Taken together, this suggests that an intra nerve AAV2/9-mediated gene therapy represents an effective and attractive therapy for myelin-related CMT diseases.

## Results

### Broad and specific transduction of mSC is reached after AAV2/9 injection in sciatic nerves of mammals

We first investigated the transduction efficiency of recombinant single-stranded AAV2/9 and AAV2/rh10 vectors expressing eGFP under a CAG promoter after injection into the sciatic nerves of mice and rats. These injections were done using a non-traumatic microinjection protocol previously described **[18]**. Briefly, fine glass needles containing a viral solution stained with Fast Green were introduced in the sciatic nerve and the solution was slowly injected using multiple short-time pressure pulses. Injections of 5 × 10^10^ vg (vector genome)/nerve and 1.8 × 10^11^ vg/nerve of AAV2/9-CAG-eGFP in adult mouse and rat respectively resulted in the transduction of a significant amount of mSC (Fig. 1a), characterized by a circular shape (Fig. 1a, inserts). At the injection site, 93% and 80% of mSC were transduced in adult mouse and adult rat respectively (Table 1). The injection of the same amount of AAV2/rh10-CAG-eGFP vector resulted in the transduction of mSC but with a lower efficiency (Fig. 1a, Table 2). Using a regular 22G syringe, we also injected 5 × 10^12^ vg/nerve of each vector into the sciatic nerve of an adult nonhuman primate (NHP). A similar high transduction rate was observed for mSC using AAV2/9-CAG-eGFP but not AAV2/rh10-CAG-eGFP (Fig. 1a, Table 1, Table 2). Similar transduction efficiencies were obtained in newborn mice and rats (Postnatal day (P) 2 and P3 and rats (P6 and P7)) when myelination starts (Fig. 1a, Table 1, Table 2).

**Table 1.** Quantification of the transduction pattern after intra-nerve injection of AAV2/9 - CAG-eGFP in rodents and NHP. Animal groups were composed of 3 animals except for NHP (one adult). Mouse and rat pups were injected at P2-P3 and P6-P7 respectively. Adult mice and rats were injected at 2-3 months old. Adult NHP was injected at 3.7 years old. All animals were sacrificed one month post-injection. The transduction rate is the percentage of transduced mSC on the overall number of mSC per section. Proximal distances from the injection site were 2, 3 and 4 cm for mice, rats and NHP respectively. Distal distances from the injection site were 0.5, 1 and 2 cm for mice, rats and NHP respectively. The specificity is the ratio of mSC, non mSC and axons transduced on the overall number of transduced cells. The results are expressed as the mean ± SD. NA, Not Available.

|  |  | *Mouse* |  | *Rat* |  | *NHP* |
| --- | --- | --- | --- | --- | --- | --- |
|  |  | *Adult* | *Pup* | *Adult* | *Pup* | *Adult* |
| Transduction rate (%) | Proximally from SI | 63 ± 24 | 74 ± 7 | NA | 81 ± 7 | 21 ± 2 |
|  | Injection Site | 93 ± 2 | 85 ± 15 | 80 ± 14 | 87 ± 1 | 68 ± 3 |
|  | Distally from SI | 91 ± 2 | 74 ± 14 | NA | 50 ± 24 | 69 ± 5 |
| Specificity (%) | mSC | 97 ± 4 | 87 ± 12 | 89 ± 9 | 95 ± 2 | NA |
|  | Non mSC | 3 ± 3 | 2 ± 2 | 7 ± 5 | 4 ± 1 | NA |
|  | Axons | 0 | 11 ± 11 | 4 ± 4 | 1 ± 1 | NA |

**Table 2.** Quantification of the transduction pattern after intra-nerve injection of AAV2/rh10-CAG-eGFP in rodents and NHP. Adult mouse and rat pup groups were composed of 3 animals. NHP adult group was composed of one adult. Adult mice, rat pups and adult NHP were respectively injected at 2-3 months old, P6-P7 and 2.3 years old. All animals were sacrificed one month post-injection. The transduction rate is the percentage of transduced mSC on the overall number of mSC per section. Proximal distances from the injection site were 2, 3 and 4 cm for mice, rats and NHP respectively. Distal distances from the injection site were 0.5, 1 and 2 cm for mice, rats and NHP respectively. The specificity is the ratio of mSC, non mSC and axons transduced on the overall number of transduced cells. The results are expressed as the mean ± SD. NA, Not Available

|  |  | *Adult mouse* | *Rat pup* | *Adult NHP* |
| --- | --- | --- | --- | --- |
| Transduction rate (%) | Proximally from SI | 42 ± 22 | 52 ± 3 | < 1 |
|  | Injection Site | 51 ± 11 | 46 ± 5 | < 1 |
|  | Distally from SI | 42 ± 16 | 5 ± 6 | < 1 |
| Specificity (%) | mSC | 82 ± 17 | 95 ± 1 | NA |
|  | Non mSC | 4 ± 4 | 4 ± 1 | NA |
|  | Axons | 14 ± 13 | 1 ± 1 | NA |

**Fig. 1.**
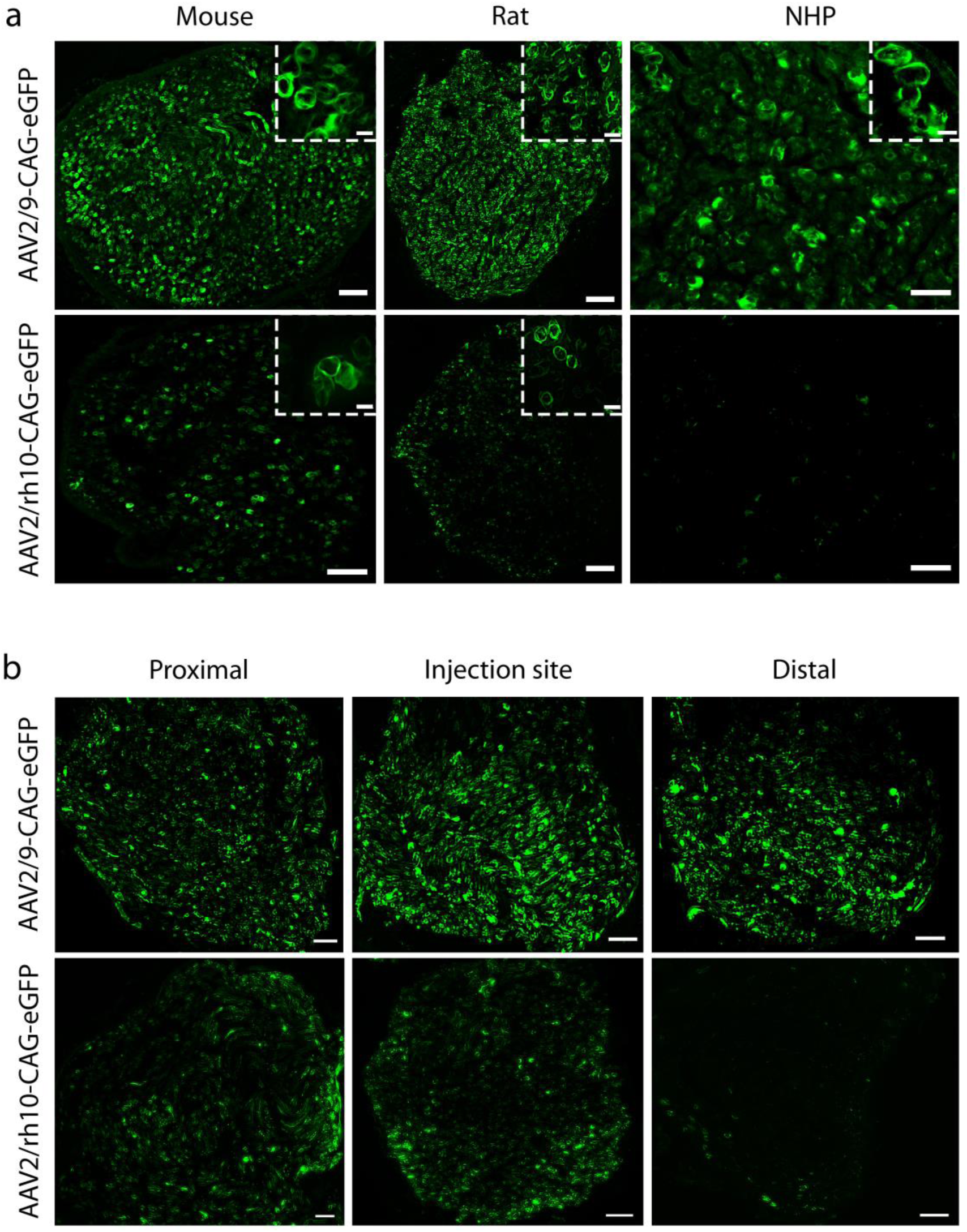
Transduction rate and diffusion after intra-nerve injection of AAV2/9-CAG-eGFP and AAV2/rh10-CAG-eGFP. Representative images of eGFP protein expression after intra-nerve injection of AAV2/9 and AAV2/rh10-CAG-eGFP in adult mouse (2-3 months old, 5 × 10^10^ vg/nerve in 8 µl, N = 3 animals per group), rat pup (P6-P7, 1 × 10^11^ vg/nerve in 8 µl, N = 3 animals per group) and adult NHP (3.7 and 2.3 years old for AAV2/9 and AAV2/rh10, 5 × 10^12^ vg/nerve in 416 µl, N = 1 animal per group). All animals were sacrificed one month post-injection. (**a**) Cross-sections of injected sciatic nerves of adult mouse, rat pup and adult NHP at the injection site. Inserts show the circular shape of transduced mSC. Scale bars: 100 µm (mouse), 50 µm (rat and NHP), 10 µm (inserts). **(b**) Cross-sections of injected sciatic nerves of rat pup at the injection site, proximally (3 cm) and distally (1 cm) from the injection site. Scale bars: 50 µm

We also evaluated the diffusion of the vectors along mouse and rat sciatic nerves collecting injected nerve samples proximally and distally from the injection point in order to cover the full length of the nerve. This diffusion was very significant for AAV2/9-CAG-eGFP as the average transduction rate was 73% at these two distant points in both mice and rats (Fig. 1b, Table 1, Table 2). In the NHP sciatic nerve injected with AAV2/9-CAG-GFP vector, the diffusion was evaluated 4 cm proximally and 2 cm distally from the injection point covering 30 to 50% of the nerve length. The transduction rate was between 21% and 69% at these distant points (Table 1). The difference resulted from the direction of the needle when inserted into the nerve.

The specificity of the viral transduction was then measured through a more extensive characterization of the transduced cells. First, we imaged adult rat sciatic nerves 12 months after injection in newborn pups using Coherent Antistokes Raman Scattering (CARS) non-linear microscopy technique. This allows myelin sheath imaging without any labeling [21]. Using this technique on an intact nerve in a longitudinal view, we observed that the majority of transduced cells were myelinated cells (Fig. 2a). In addition, typical morphological characteristics of mSC were seen through eGFP labeling such as Schmidt-Lanterman incisures (Fig. 2a, arrows), Cajal’s bands (Fig. 2a, arrowheads) and nodes of Ranvier (Fig. 2a, stars). Sciatic nerve cross-sections were immunostained for axonal and myelin markers allowing to definitively identify mSC as eGFP-labelled myelin-positive cells surrounding axons (Fig. 2b, Table 1). As some non-myelinating cells (non-mSC) located in the nerve, such as unmyelinated Schwann cells and fibroblasts, are not easily seen in cross-sections, we teased sciatic nerve fibers on a glass slide and counted eGFP-positive mSC, non-mSC and axons (Fig. 2c). On 100 cells transduced with AAV2/9-CAG-eGFP vector after intra-nerve injection in rat pups, 95 were mSC, 4 non-mSC and 1 was an axon (Table 1), showing that AAV2/9 is highly specific for mSC when injected into the nerve. The same specificity was observed after intra-nerve injection in adult rats, adult mice and mouse pups (Table 1, Table 2). Taken together the data highlight the high transduction rate, diffusion and specificity of AAV2/9 for mSC, as well as the long term expression of the transgene after intra-nerve injection.

**Fig. 2.**
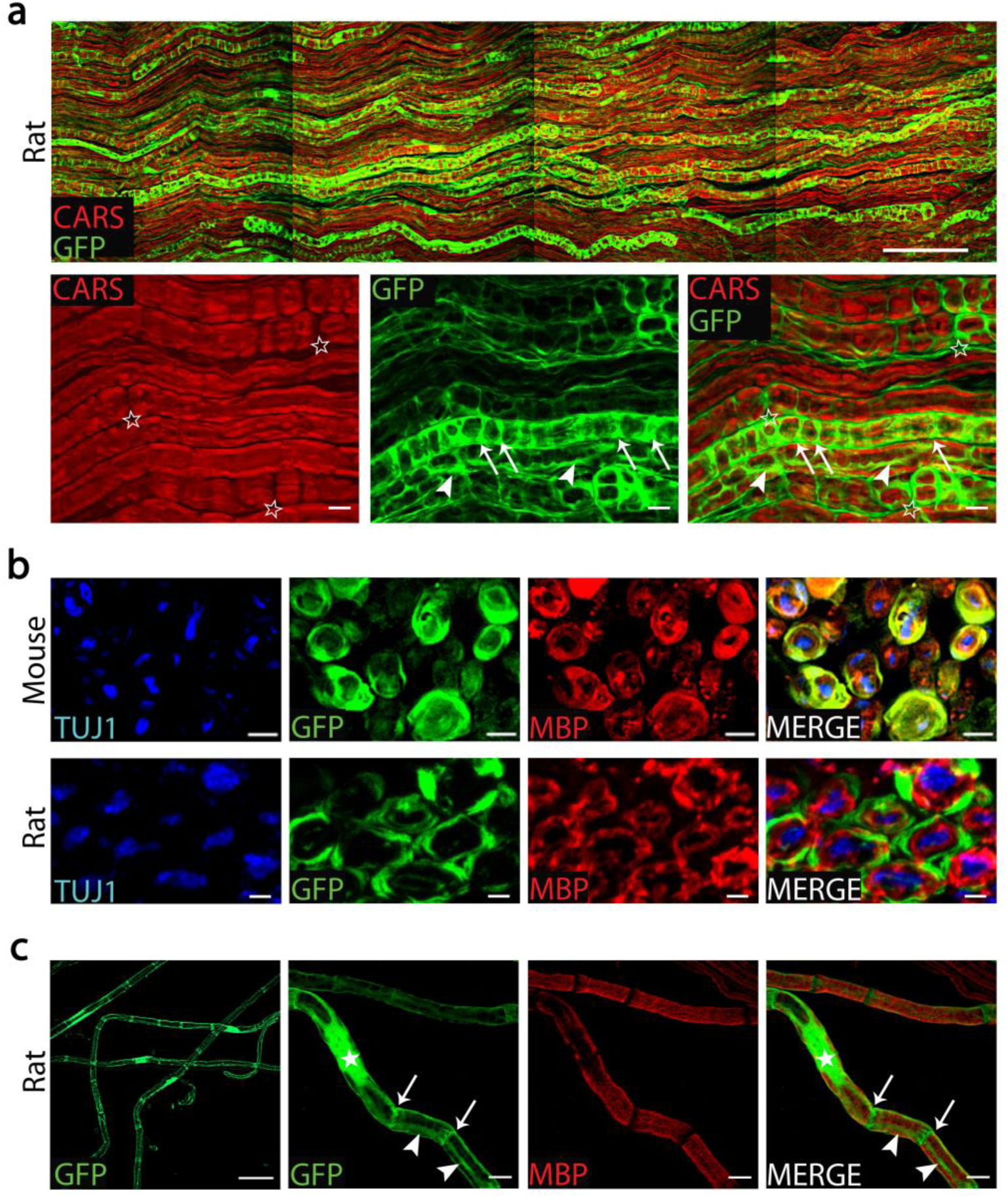
AAV2/9 transduction is specific for mSC after intra-nerve injection. (**a**) Upper panel: CARS (red) and eGFP (green) imaging of a 1cm-long sciatic nerve injected with AAV2/9 expressing eGFP in rat pup (1 × 10^11^ vg/nerve in 8 µl) 12 months post-injection. Scale bar: 100µm. Lower panel: Higher magnifications showing specific features of transduced mSC, such as Schmidt-Lanterman incisures (arrows), Cajal’s band (arrowheads) and nodes of Ranvier (stars). Scale bar: 10 µm. (**b**) Representative images of immunostainings for myelin MBP (red) and axonal Tuj1 (blue) on sciatic nerve cross-sections of adult mouse (injection 2-3 months old, 5 × 10^10^ vg/nerve in 8 µl, N=3 animals) and of rat pup (injection P6-P7, 1 × 10^11^ vg/nerve in 8 µl, N = 3 animals) injected with AAV2/9-CAG-eGFP show that mSCs expressing eGFP (green) partially colocalize with MBP and surround axons. Animals were sacrificed one month post-injection. Scale bar: 4 µm (mouse) and 2 µm (rat). (**c**) Left panel: Representative images of teased fibers from a rat sciatic nerve injected with AAV2/9-CAG-eGFP (injection P6-P7, 1 × 10^11^ vg/nerve in 8 µl, N = 3 animals) and sacrificed one month post-injection. eGFP is expressed in several mSC characterized by a long tubular shape. Scale bar: 30 µm. Right panels: higher magnification showing the immunostaining for myelin MBP (red) of teased fibers. eGFP (green) is expressed in the nucleus (star), the Cajal’s bands (arrowheads) and the Schmidt-Lanterman incisures (arrows) of mSCs. Scale bar: 10 µm

### AAV2/9 expressing PMP22 shRNAs restore wild-type PMP22 level in mutant CMT1A rats

The high and specific transduction of mSC in mouse, rat and NHP by AAV2/9 lead us to evaluate its use as a vector to carry a therapeutic tool into defective mSC in CMT1 diseases. As a proof of concept, we chose a rat model of CMT1A in which mouse PMP22 is overexpressed [57]. This model closely mimics the human disease: immediately after peripheral nerve myelination begins, nerves show hypomyelination, shorter internodal lengths, deficit in large fibers, hypermyelination of small fibers, myelin sheath defects and demyelination [13, 15, 16]. We designed a therapeutic vector carrying both a small shRNA targeting mouse PMP22 mRNA under a U6 promoter and a CAG-eGFP reporter cassette. Two effective shRNAs (sh1 and sh2) were characterized *in vitro* on mouse MSC80 and rat RT4-D6P2T Schwann cell lines. Both shRNAs significantly downregulated mouse PMP22 level in a dose-dependent manner, while only sh1 was effective on rat PMP22 level (Fig. 3).

**Fig. 3.**
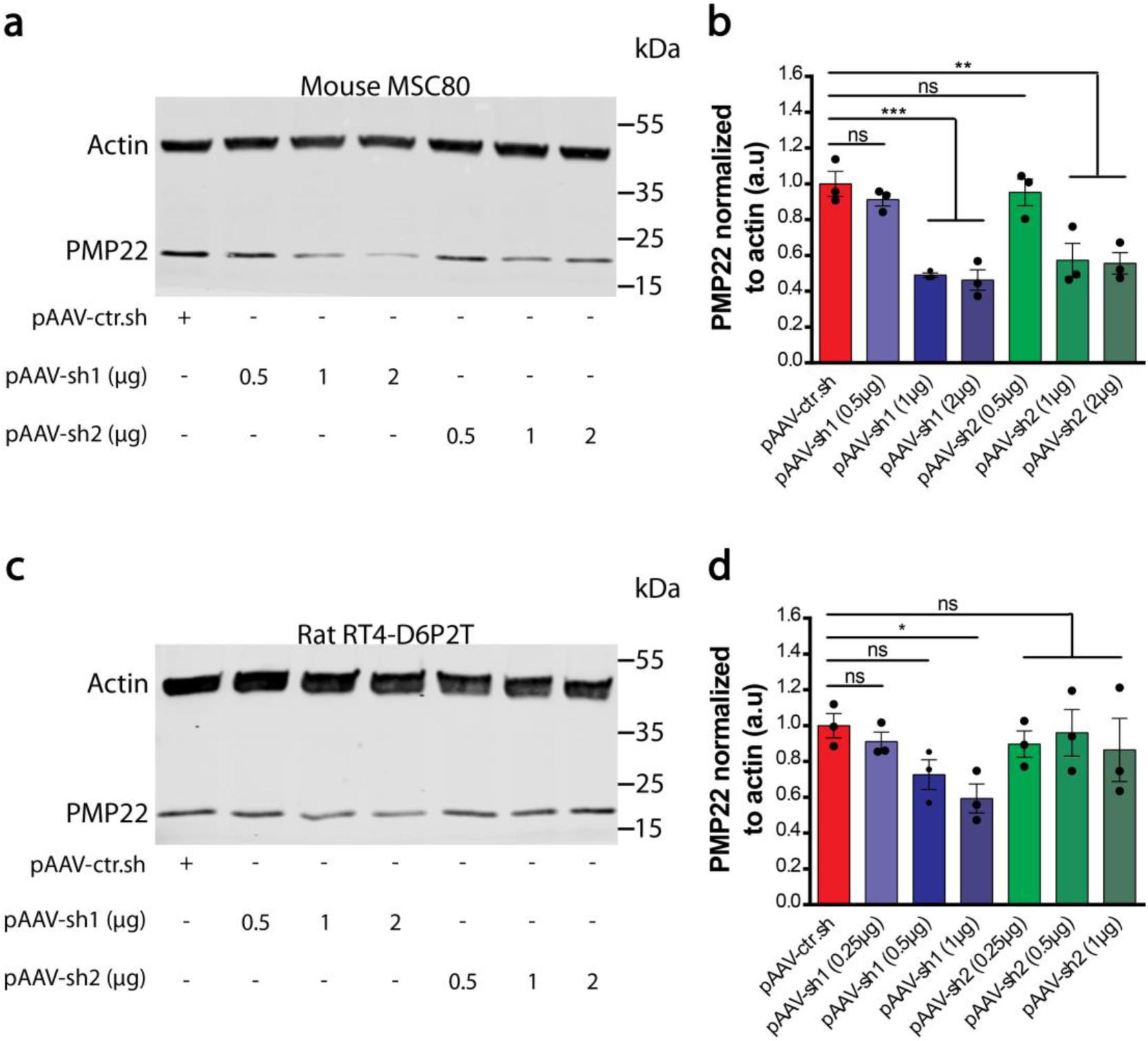
sh1 and sh2 efficiently silence mouse PMP22 while only sh1 silences rat PMP22. (**a**) Western blot analysis of PMP22 protein in mouse MSC80 Schwann cell line transfected with pAAV-sh1, pAAV-sh2 or pAAV-ctr.sh at the indicated dose per well (µg of DNA). (**b**) Graph shows mean mouse PMP22 level in transfected MSC80 lysates normalized to actin (N = 3 per group). (**c**) Western blot analysis of PMP22 protein in rat RT4-D6P2T Schwann cell line transfected with pAAV-sh1, pAAV-sh2 or pAAV-ctr.sh at the indicated dose per well (µg of DNA). (**d**) Graph shows mean rat PMP22 level in transfected RT4-D6P2T lysates normalized to actin (N = 3 per group). Statistical tests show one-way ANOVA followed by Dunnett’s post hoc test for pAAV-sh1 and pAAV-sh2. * *P* < 0.05, ** *P* < 0.01, *** *P* < 0.001; ns, not significant. a.u, arbitrary unit. All error bars show SEM.

AAV2/9 vectors expressing sh1 or sh2 (AAV2/9-sh1 and AAV2/9-sh2) were then injected bilaterally (1 × 10^11^ vg/nerve) in the sciatic nerves of CMT1A rat pups (P6-P7) (groups CMT1A sh1 and CMT1A sh2 respectively). WT littermates and CMT1A rat pups injected with AAV2/9 carrying a control shRNA (AAV2/9-ctr.sh) were used as controls (groups WT ctr.sh and CMT1A ctr.sh respectively). Treated and control animals were followed for as long as 12 months after injection.

First, we investigated the efficiency of both AAV2/9-sh1 and AAV2/9-sh2 vectors to reduce PMP22 expression in the nerves. As PMP22 is expressed in the myelin and myelin is altered in CMT1A, we normalized PMP22 expression to expression level of Myelin Protein Zero (MPZ). While PMP22 level was increased in CMT1A ctr.sh compared to WT ctr.sh, transduction of mSC by AAV2/9-sh1 and AAV2/9-sh2 decreased PMP22 level back to control WT ctr.sh (Fig. 4a and b). This indicated that AAV2/9 vectors carrying shRNA targeting PMP22 were able to correct PMP22 overexpression in CMT1A rats. No downregulation beyond that of control levels was observed in CMT1A sh1 and CMT1A sh2 animals.

**Fig. 4.**
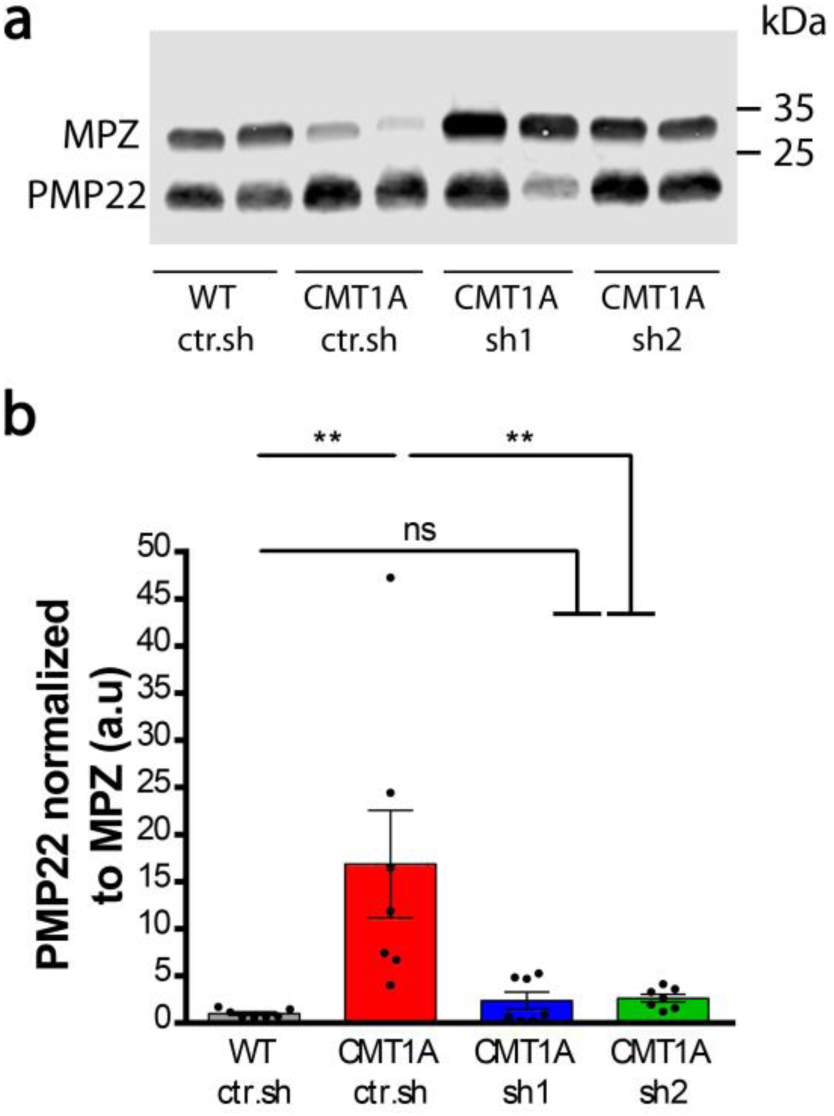
Intra-nerve injection of AAV2/9-sh1 and sh2 corrects PMP22 overexpression in CMT1A sciatic nerves. (**a**) Representative image of Western blot showing PMP22 and MPZ protein levels in rat sciatic nerve lysates from WT ctr.sh, CMT1A ctr.sh, CMT1A sh1 or CMT1A sh2 3 months after injection. (**b**) Graph shows mean PMP22 level in sciatic nerve lysates normalized to MPZ level (N = 7 animals per group). Statistical test shows one-way ANOVA followed by Tukey’s post hoc. ** *P* < 0.01; ns, not significant; a.u, arbitrary unit. All error bars show SEM

### AAV2/9-PMP22 shRNAs reduce myelinated fibers defects in the sciatic nerve of CMT1A rats

As CMT1A disease affects mSC and leads to the loss of myelin, we assessed the amount of myelin marker MPZ after AAV2/9-sh1 and -sh2 treatment using Western blot (Fig. 5a). While CMT1A rats showed a lower level of MPZ than WT ctr.sh rats, MPZ levels in CMT1A sh1 and sh2 rats were not statistically different from WT ctr.sh rats (Fig. 5b).

**Fig. 5.**
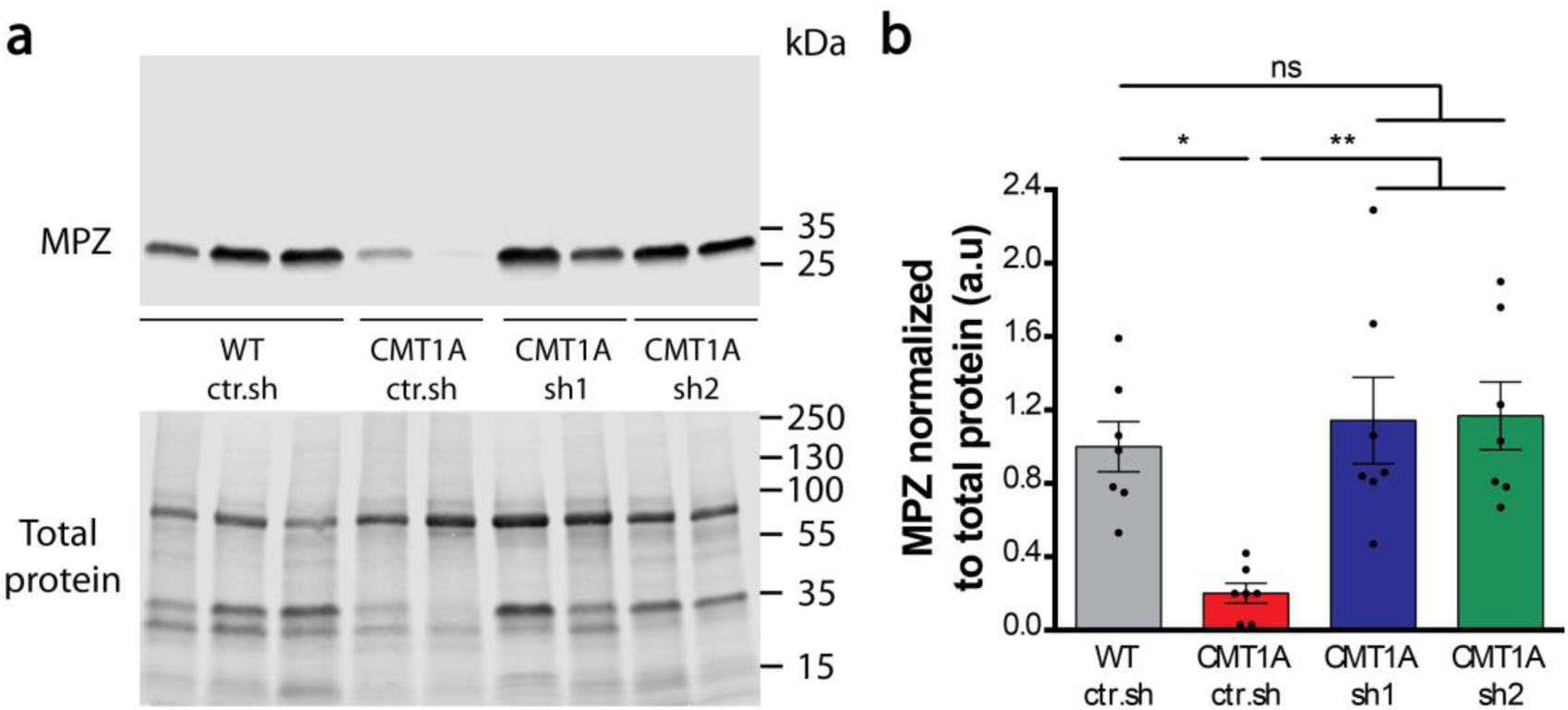
Intra-nerve injection of AAV2/9-sh1 and sh2 in CMT1A rats normalizes MPZ expression back to WT rat level. (**a**) Representative images of Western blot showing MPZ protein (upper panel) and total protein (lower panel) levels in sciatic nerve lysates from WT ctr.sh, CMT1A ctr.sh, CMT1A sh1 or CMT1A sh2 3 months after injection. (**b**) Graph shows mean MPZ level in sciatic nerve lysates normalized to total protein level per nerve (N = 7 animals per group). Statistical test shows one-way ANOVA followed by Tukey’s post hoc. * *P* < 0.05 ** *P* < 0.01; ns, not significant. a.u, arbitrary unit. All error bars show SEM

Second, we focused on the specific myelin defects observed in the CMT1A rat model using CARS imaging on freshly fixed nerves. Indeed, in a previous study, we had characterized CMT1A rat myelin sheath defects using this imaging technique [21]. The CMT1A myelin sheath showed a high heterogeneity compared to WT rat littermates: thin myelin sheath, demyelination (Fig. 6a, arrows), followed by focal hypermyelination (Fig. 6a, arrowheads), and myelin degeneration with ovoids formation (Fig. 6a, stars). While still present, all these defective features were less abundant in CMT1A rats treated with AAV2/9-sh1 or –sh2 (Fig. 6a). A typical feature of CMT1A nerves is the shorter internodal distance [13, 54]. Using CARS, we found that the number of nodes of Ranvier per fiber increased in CMT1A ctr.sh nerves compared to WT ctr.sh nerves (Fig. 6b and c), indicating shorter internodes. When CMT1A rats were treated with AAV2/9-sh1 or -sh2, node number per fiber decreased significantly, showing that internodes were longer.

**Fig. 6.**
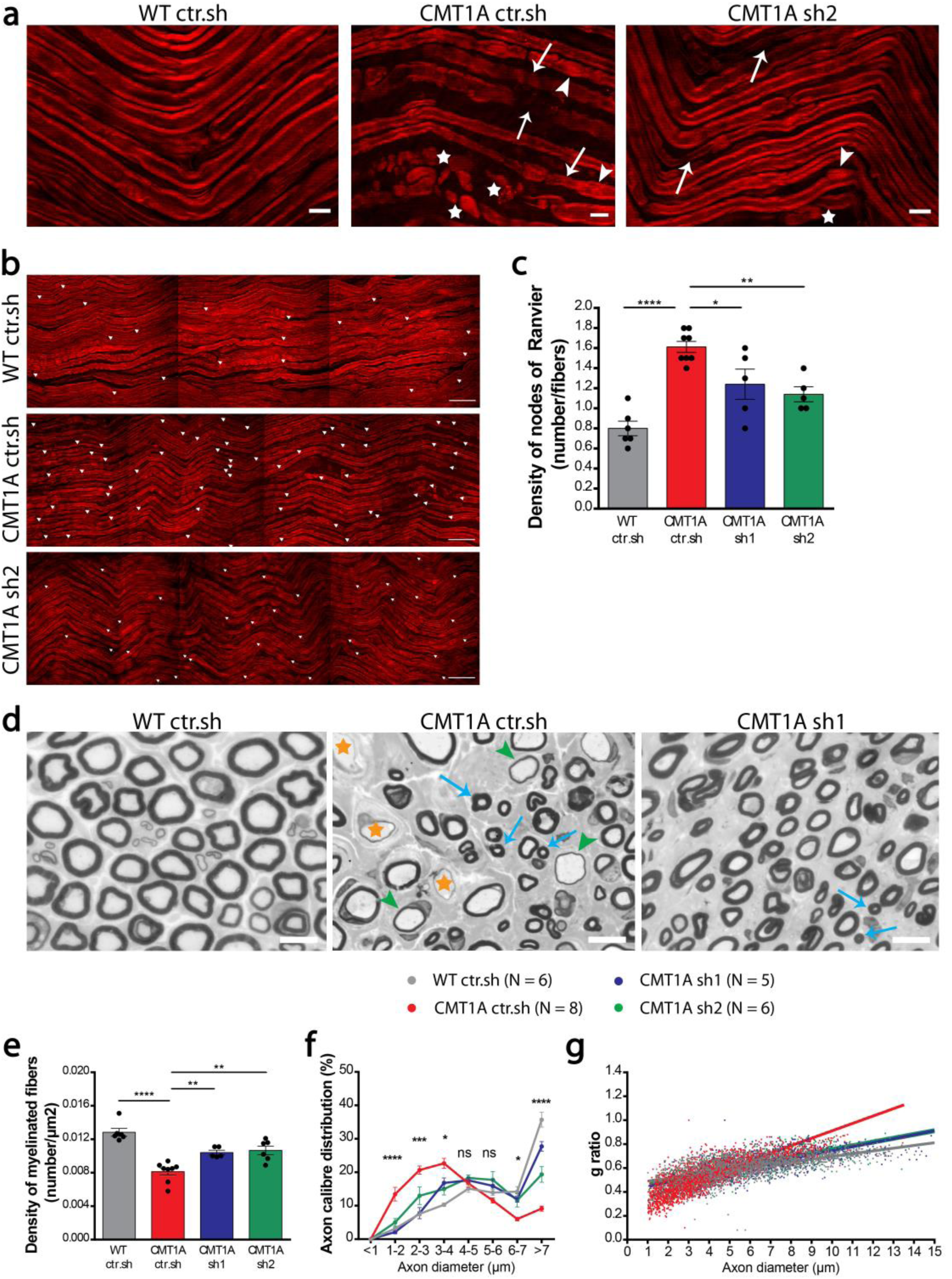
Intra-nerve injection of AAV2/9-sh1 and sh2 corrects the myelin sheath defects in CMT1A rats. **(a** and **b**) Representative images of CARS imaging on WT ctr.sh, CMT1A ctr.sh and CMT1A sh2 rat sciatic nerves 12 months post-injection. (**a**) CMT1A ctr.sh nerve shows typical myelin sheath defects such as thin myelin sheath (arrows), focal hypermyelination (arrowheads) and myelin degeneration with myelin ovoids (stars). These defects are less abundant in CMT1A sh2 rat sciatic nerves. Scale bars: 10 µm. (**b**) Nodes of Ranvier were labelled with arrowheads. Nodes of Ranvier are more abundant in CMT1A ctr.sh nerve. Scale bars: 60 µm. (**c**) Graph showing the mean number of nodes of Ranvier on the total number of myelinated fibers per field (N = 5 to 8 animals). Statistical test shows one-way ANOVA followed by Tukey’s post hoc. * *P* < 0.05, ** *P* < 0.01; *** *P* < 0.001; ns, not significant. (**d**) Representative images of electron microscopy semi thin sections on WT ctr.sh, CMT1A ctr.sh and CMT1A sh1 rat sciatic nerves 12 months post-injection. CMT1A ctr.sh nerve shows typical myelinated fiber defects such as large demyelinated axons (orange stars), large hypomyelinated axons (green arrowheads) and small hypermyelinated axons (blue arrows). These defects are less abundant in CMT1A sh1 rat sciatic nerve. Scale bars: 10 µm. Graphs showing (**e**) the mean number of myelinated fibers per area unit (µm^2^), (**f**) the mean percentage of axon caliber distribution per axon diameter, and (**g**) g-ratio relative to axon diameter (N = 5 to 8 animals per group). Statistical tests show one-way ANOVA followed by Tukey’s post hoc (**c** and **e**) or two-way ANOVA followed by Dunnett’s post hoc (**f**). * *P* < 0.05, ** *P* < 0.01; *** *P* < 0.001, **** *P* < 0.0001; ns, not significant. All error bars represent SEM.

Finally, we investigated the myelin sheath structure using electron microscopy semi-thin nerve sections. Light microscopy imaging of these sections showed typical features of rat CMT1A disease: lower density of myelinated fibers, hypomyelination or demyelination of large axons and hypermyelination of small axons (Fig 6d). These features were still present in the nerve of treated animals but at lesser extent (Fig 6d). We found that AAV2/9-sh1 and –sh2 treatment significantly increased the myelinated fibers density (Fig. 6e), the number of large myelinated axons and proportionally reduced the number small myelinated axons (Fig. 6f). Finally, the g-ratio (axon diameter/myelinated fiber diameter) was increased for small caliber axons and decreased for large caliber axons, not significantly different to control WT ctr.sh values (Fig 6g). Taking together, these data indicate that the treatment with AAV2/9-sh1 or –sh2 prevents myelin loss and the occurrence of myelinated fibre defects in CMT1A rats.

### PMP22 silencing prevents motor and sensitive defects on the long term in CMT1A rats

As CMT1A is a myelin-related disease, one of the first symptoms to occur is the decrease of the nerve conduction velocity (NCV). Accordingly, NCV was significantly reduced in CMT1A ctr.sh animals as soon as one month after birth compared to control WT ctr.sh animals (Fig. 7a). When CMT1A animals were treated with AAV2/9-sh1 or-sh2, the NCV remained not significantly different to WT ctr.sh values at all-time points for at least 12 months (Fig. 7a).

**Fig. 7.**
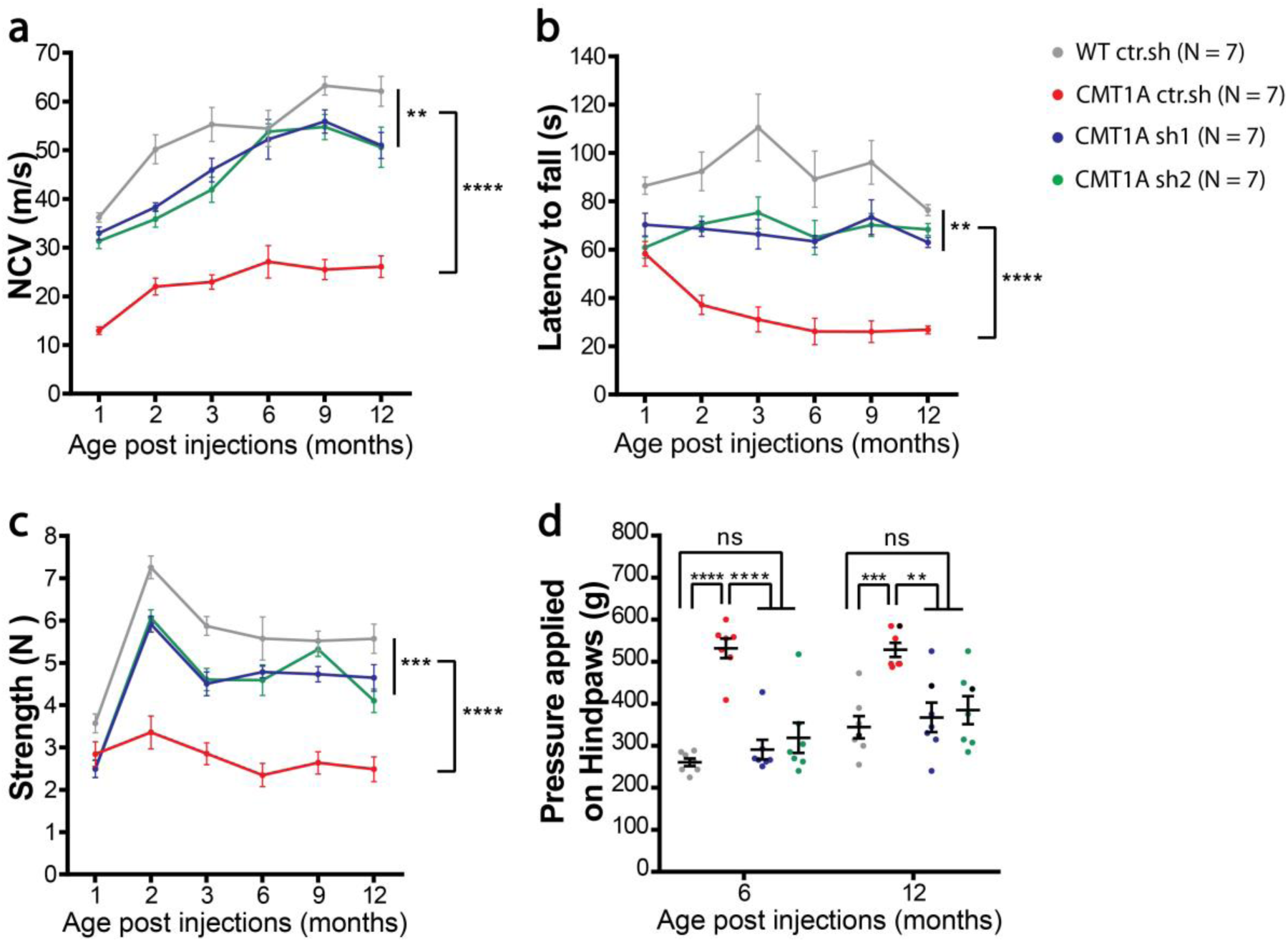
Intra-nerve injection of AAV2/9-sh1 and sh2 prevents motor and sensory defects on the long term in CMT1A rats. Graphs showing (**a**) NCV (meter/second), (**b**) Rotarod test (second), (**c**) grip test (Newton) 1 to 12 months after injection and (**d**) Randall Selitto test (gram) 6 and 12 months after injection in WT ctr.sh, CMT1A ctr.sh, CMT1A sh1 and CMT1A sh2 animals (N = 7 animals per group). Statistical analysis shows two-way ANOVA followed by Tukey’s post hoc (**a, b** and **c**) or one-way ANOVA followed by Tukey’s post hoc for 6 and 12 months post-injection (**d**). ** *P* < 0.01; *** *P* < 0.001, **** *P* < 0.0001; ns, not significant. Results are expressed as mean ± SEM.

The strong decrease of NCV in diseased animals resulted in a clumsy behavior when animals crossed a narrow beam and the treatment prevented this defect (Online Resource 1 and 2). Motor behavior tests of rotarod and grip strength also showed reduced performances of CMT1A ctr.sh animals compared to control WT ctr.sh animals, starting 2 months after birth and these deficiencies were largely prevented by the treatment with AAV2/9-sh1 and – sh2 (Fig. 7b and c). No significant evidence of a reduction in the effectiveness of the treatment could be observed even 12 months after the treatment.

CMT1A disease shows a reduced sensitive perception due to both a defect in myelinated sensory fibers and a loss of sensory axons on the long term [31]. Consequently, we performed Randall-Selitto test to assess the effects of the therapeutic vectors on the mechanical pain sensitivity of lower limbs at 6 months and 12 months post-injection (Fig. 7d). We first observed an increase in the mechanical nociceptive threshold of CMT1A ctr.sh animals as compared to the WT ctr.sh group indicating hypoalgesia. Treatment with both AAV2/9-sh1 and –sh2 completely prevented this sensitive deficit (Fig. 7d). Consequently, treatment with AAV2/9 vectors that reduces PMP22 levels in mSC constitutes an efficient and long-term preventive treatment for CMT1A symptoms in rats.

### Human skin biomarkers reliably and robustly discriminate treated from sham and wild-type animals

Recent clinical trials for drugs targeting CMT diseases have illustrated the lack of outcome measures sensitive enough for such chronic peripheral neuropathies [43, 63]. We therefore tested whether recently discovered human skin mRNA biomarkers [14] could be used as an outcome measure for our gene therapy efficiency. Forepaw glaber skin was collected at 12 months in all cohorts and the expression of nine identified biomarkers was quantified using RT-qPCR. Some of these biomarker expressions were significantly differentially expressed in the skin of CMT1A sh1 and CMT1A sh2 animals versus CMT1A ctr.sh animals (Online Resource 3). However, the expression levels remained highly heterogeneous among animals, reflecting the variability of CMT1A disease expression that is recapitulated in transgenic rats [15]. We therefore performed a multi-variated principle component analysis (PCA), including the entire battery of nine validated human biomarkers. In this analysis, biomarkers allowed a significant segregation of WT ctr.sh animals from CMT1A ctr.sh animals, and of CMT1A sh1 and CMT1A sh2 animals from CMT1A ctr.sh animals (Fig. 8a). These data indicated that biomarkers expression in the paw skin allows for detecting the outcome of the gene therapy treatment in CMT1A rats.

**Fig. 8.**
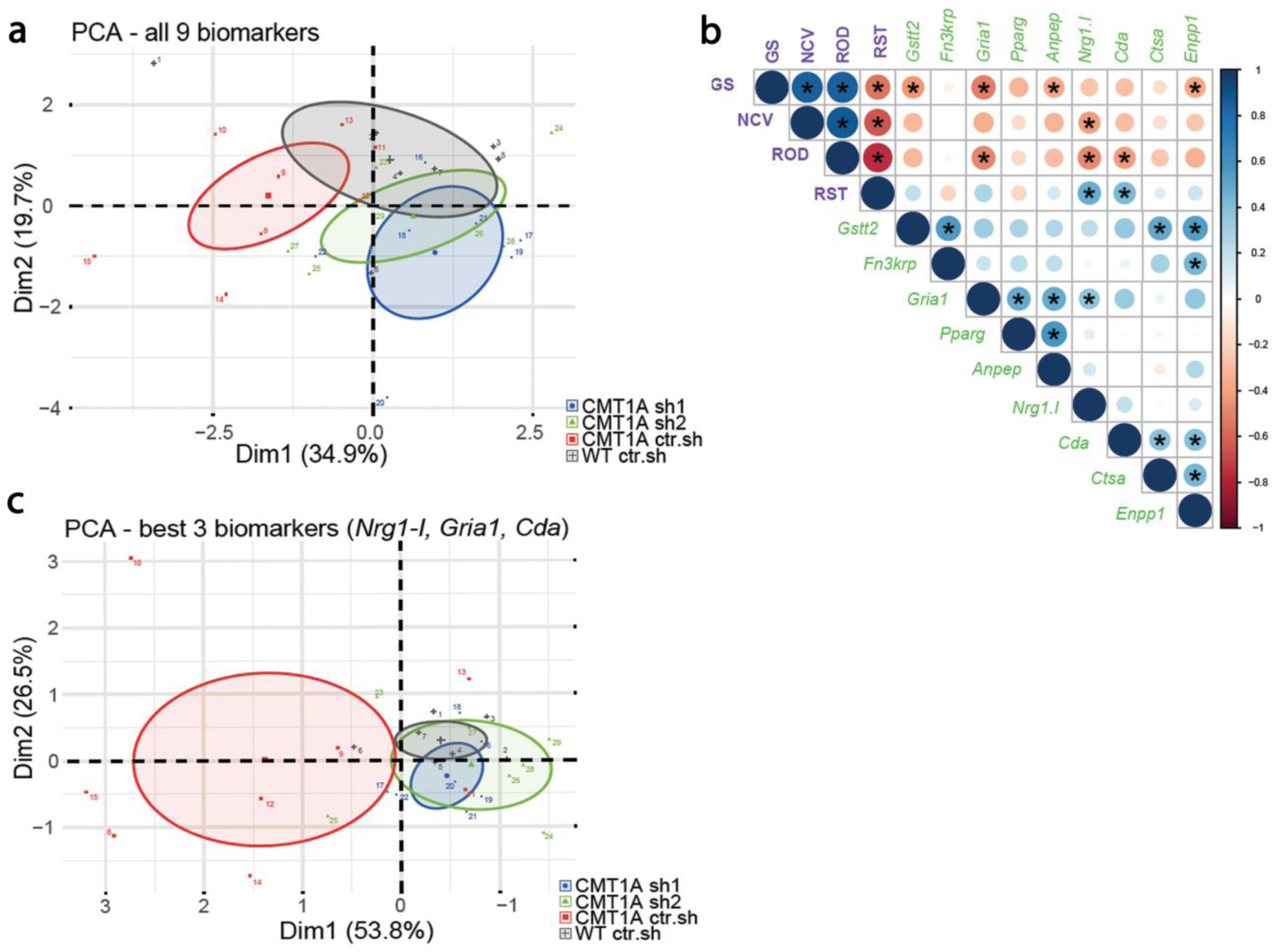
Multi-variated analysis of skin biomarkers and sensory-motor phenotypes allows the detection of the therapy outcome. (**a**) Principal component analyses (PCA) of all nine transcriptional biomarkers in forepaw skin biopsies on 12 months old animals. Note little to no overlap between WT ctr.sh (grey, n = 7) and CMT1A ctr.sh (red, n = 8), whereas the treated groups, CMT1A sh1 (blue, n = 7) and CMT1A sh2 (green, n = 7), show more overlap with the WT ctr.sh group than with the CMT1A ctr.sh group. The mean of each group is given as a center point including the confidence interval (95%) given as an ellipse. (**b**) Correlation matrix from all animals (total n = 28 with n=7 per group) including the expression levels of the skin biomarkers (green labels) and the four functional phenotypic analyses (purple labels): GS, grip strength; NCV, nerve conduction velocity; ROD, Rotarod; RST, Randall-Selitto test). Shown is data from a Pearson’s correlation analyses with graphical representation of the correlation coefficients, from red (−1) to blue (+1), and the respective p values (lower with increased circle size, asterisks indicate p<0.05). (**c**) Principal component analyses (PCA) of the three best biomarkers (*Nrg1-I, Gria1, Cda*; see correlation matrix in (**b**)) in forepaw skin biopsies on 12 months old animals (same analysis as in (**a**)).

To go further, we performed a correlation analysis including both functional phenotypes (Rotarod-ROD, grip strength test-GS, Nerve Conduction Velocity-NCV and Randall-Selitto test-RST) and biomarkers expression at 12 months. While correlations were more significant (stars and large disk size) within the phenotypic test group or more numerous within the biomarkers group (11/27), significant correlations occurred between functional phenotypes and biomarkers groups (10/27) (Fig. 8b). *Nrg1.1* (3 correlations), *Gria1* (2), *Cda* (2), *Gstt2* (1) *Anpep* (1) and *Enpp1* (1) were the genes whose skin expression correlated the best with the phenotype improvement of CMT1A rats following gene therapy. Indeed, when the 3 most relevant biomarkers (*Nrg1.1, Gria1* and *Cda*) were selected for the PCA analysis, the segregation between CMT1A ctr.sh and CMT1A sh1 and CMT1A sh2 animals was clear (Fig. 8c). According to the correlation matrix, the grip strength test (4 correlations) was slightly more relevant than Rotarod (3), Randall-Selitto (2) and NCV (1) tests. However, Rotarod (3) and Randall-Selitto (2) tests were sufficient to robustly cover the 3 most relevant biomarkers (Fig. 8b). Taken together these data indicate that a combinative score grouping functional tests and skin biomarkers analysis constitute a reliable and robust measure of CMT1A gene therapy outcome in CMT1A rats.

### Off-target transduction and immune response to the vector are limited after intra-nerve injections

Regarding a future clinical trial for this gene therapy approach, we investigated the biodistribution of the AAV2/9 vector after its injection in the sciatic nerve of rat pups 3 months post-injection. Tissues were collected in AAV2/9 injected animals and vector genome copy numbers were analyzed using normalized and standardized qPCR (Table 3). Among the 32 analyzed animals, 29 showed vector genome copies in the sciatic nerve indicating that the injection technique was highly reliable with a success rate of 91 %. Among all groups, the average amount of vector genome relative to diploid genome (vg/dg) in the sciatic nerve was 0.54 ± 0.09 (Table 3). Four out of 32 animals displayed very low levels of vector genome copies in the liver (0.005 ± 0.003 vg/dg), three in the heart (0.004 ± 0.001 vg/dg) and five in lumbar dorsal root ganglia L4 L5 (0.004 ± 0.001 vg/dg). One out of 7 animal displayed vector genome copy in the blood (0.005 vg/dg). No vector genome copy was detected in other tested organs (brainstem, spinal cord, kidney, spleen).

**Table 3.** AAV2/9 biodistribution in sciatic nerves and different organs 3 months after intra-nerve injection. Rats were injected at P6 -7 and sacrificed at 3 months later. The first column shows the number of positive samples (> LOQ) over all tested samples. The second column shows the transduction rate expressed in vector genome/diploid genome (vg/dg). Results are expressed as the mean ± SD. NA, Not Applicable

|  | *WT ctr.sh* |  | *CMT1Actr.sh* |  | *CMT1Ash1* |  | *CMT1Ash2* |  | *Overall* |  |
| --- | --- | --- | --- | --- | --- | --- | --- | --- | --- | --- |
|  | *Positive*<br>*/total* | *vg/dg* | *Positive*<br>*/total* | *vg/dg* | *Positive*<br>*/total* | *vg/dg* | *Positive*<br>*/total* | *vg/dg* | *Positive*<br>*/total* | *vg/dg* |
| Sciatic nerve | 8/8 | 0.40 ± 0.31 | 7/8 | 0.21 ± 0.11 | 7/8 | 0.55 ± 0.22 | 7/8 | 1.03 ± 0.86 | 29/32 | 0.54 ± 0.054 |
| Drg L4-L5 | 2/8 | 0.003 ± 0.001 | 2/8 | 0.006 ± 0.004 | 1/8 | 0.003 | 0/8 | NA | 5/32 | 0.004 ± 0.002 |
| Lumbar spinal cord | 0/8 | NA | 0/4 | NA | 0/2 | NA | 0/2 | NA | 0/16 | NA |
| Heart | 2/8 | 0.004 ± 0.001 | 0/8 | NA | 1/8 | 0.003 | 0/8 | NA | 3/32 | 0.004 ± 0.001 |
| Liver | 1/8 | 0.003 | 1/8 | 0.008 | 2/8 | 0.004 ± 0.001 | 0/8 | NA | 4/32 | 0.005 ± 0.003 |
| Spleen | 0/4 | NA | 0/2 | NA | 0/1 | NA | 0/1 | NA | 0/8 | NA |
| Kidney | 0/4 | NA | 0/2 | NA | 0/1 | NA | 0/1 | NA | 0/8 | NA |
| Brainstem | 0/4 | NA | 0/2 | NA | 0/1 | NA | 0/1 | NA | 0/8 | NA |
| Whole blood | 0/3 | NA | 1/2 | 0.005 | 0/1 | NA | 0/1 | NA | 1/7 | 0.005 |

This very limited distribution of the vector prompted us to measure the immune response against the vector capsid. A validated and standardized ELISA approach was used to measure AAV2/9 neutralizing factors in sera of ten injected animals (5 WT ctr.sh and 5 CMT1A ctr.sh) and four non-injected controls (2 WT and 2 CMT1A rats). Among these samples, only two showed neutralizing factors at low titer (1/500) (Table 4). Taken together these data show that intra-nerve injections of AAV2/9 vector results in a restricted transduction of the injected nerves with limited off-target tissues and a very weak humoral immune response toward the vector.

**Table 4.** AAV2/9 neutralizing factors titration 3 months after intra-nerve injection. The first column shows the number of positive sera over all tested samples. The second column shows the titer of positive sera

|  | *Positive sera/Total* | *Titer* |
| --- | --- | --- |
| WT ctr.sh | 1/5 | 1/500 |
| CMT1Actr.sh | 1/5 | 1/500 |

## Discussion

CMT1A is the main hereditary peripheral neuropathy representing more than 50% of all these significantly disabling diseases [62]. Since 2004, several therapeutic strategies have been proposed and tested preclinically and clinically but as of yet no treatment is available for this disease [44, 45]. At the present time, the most advanced strategy is a pharmacological treatment that reached the clinical phase III (NCT02579759). Another pharmacological treatment is reaching the clinical phase I (NCT03610334) and several others are in preclinical phases [12, 16, 34, 37, 51, 58, 66]. As all these pharmacological treatments require a regular and permanent treatment with potential side effects on the long term, gene therapy represents an attractive alternative. Indeed, an indirect gene therapy approach consists of the transduction of muscle cells to increase their production of neurotrophin 3 in order to promote axon survival is in clinical trial phase I/IIa (NCT03520751) [50, 65]. Here we investigated the conditions for a successful gene therapy approach directly targeting mSC, the defective cells in the disease, through an intra nerve delivery.

Our first goal was to evaluate the transduction efficiency and the specificity of AAV2/9 and 2/rh10 serotypes, which are the main serotypes used to transduce the nervous system, regarding mSC when injected directly in the nerve. In 2015, Tanguy *et al*. [60] observed a cell transduction in the sciatic nerve of mice intravenously injected with AAV2/9 one day after birth but no detail on the cell type was provided. Hoyng *et al*. described in 2015 the transduction efficiency of AAV2/1 to 9 in rat and human Schwann cells *in vitro* and in sectioned mouse nerve segments undergoing demyelination *ex vivo* [26]. As the existing data was inconclusive, we tested AAV2/9 and AAV2/rh10 serotypes in mouse, rat and NHP *in vivo* after intra-nerve injections. We found that both serotypes were able to transduce mSCs at a high rate in all these species and at different ages (newborn and adult). AAV2/9 was significantly more efficient than AAV2/rh10 regarding the transduction rate at the injection site (83% vs 32% respectively on average). This high transduction rate is clearly correlated to the injection protocol as an intrathecal injection of the same vectors of newborn or adult mice resulted in the transduction of a large amount of neurons and glial cells [4, 22, 23, 53] but no mSC in sciatic nerves. While we cannot rule out a transduction of mSC in nerve roots close to the spinal cord after intrathecal injection, intra nerve injection appears as the most efficient way to transduce these cells *in vivo*.

The AAV2/9 specificity for mSC was found to be slightly superior to that of AAV2/rh10 serotype in these conditions (on average 92% vs 88% respectively of transduced cells are mSC). As very few axons were found positive for eGFP after AAV2/9 injection in the nerve, further restricting the transgene expression in mSC through a specific promoter appeared redundant.

We found that the intra-nerve injection of 1 × 10^11^ vg AAV2/9 in the rat pups sciatic nerve resulted in an average of 0.54vg/dg. As AAV2/9 is 95% specific for mSC in our conditions, 87% of mSC are transduced and knowing that mSC represent 51% of the cells in a sciatic nerve [59], this indicates that nearly all mSC of the nerve are transduced with one vector copy. This is consistent with the high transduction rate of mSC we observed using immunolabelling. This also indicates that the vector dose we used to transduce all mSC of the rat pups is optimal. Moreover, the high transduction rate we obtained in the other species and in particular in NHP, suggests that we also reached the optimal dose in these conditions. Further optimizations of the treatment are therefore to be found in the extent of the vector diffusion along the nerve.

We therefore evaluated the diffusion of the vectors when injected in the nerve. Both vectors diffused similarly (at least over 2.5 cm) in the adult mouse sciatic nerve. However, 5 × 10^10^ vg of AAV2/9 distributed in 8 µl was able to transduce on average 83% of mSC along 2.5 cm-length of the entire sciatic nerve, while AAV2/rh10 only transduced on average 45% of mSCs along the same length in the same conditions. This large diffusion of AAV2/9 vector was significantly superior to that of AAV2/8 vector reported by Homs *et al.* (2.5 cm for AAV2/9 vs 0.8 cm for AAV2/8) [25]. Similar results were obtained for AAV2/9 in the rat pup nerve (1 × 10^11^ vg in 8 µl transduced on average 73% of mSC over 4 cm) and in NHP nerve (5 × 10^12^ vg in 416 µl transduced on average 53% of mSC over 6 cm). This represented the total length of rat pups and adult mice sciatic nerves and 30-50% of the adult NHP sciatic nerve. The injection technique we set up to inject viral vectors in peripheral nerves of adult mice and rat pups [18] is essential in this significant diffusion ability. Indeed, injections are done with a very fine needle under 1.5 to 2 bars pressure by successive small pulses of a few nanoliters each. The fine needle allows for perforating the sheath that surrounds the fibers (epineurium and perineurium) while preventing injury of these fibers. The short and moderated pressure pulses allow injecting a relatively large volume without inducing pressure injury in the nerve. In addition, it prevents the leak of the viral solution out of the nerve due to the immediate tissue resistance to large volume increase. Adapting and using this technique for NHP intra nerve injection will most likely increase the spread of the vector in this species also. In patients, while transdermal intra-nerve injections are not uncommon in particular during locoregional anesthesia [28], the direct injection in the nerve is presently undesired because of the toxic nature of concentrated anesthetics and the risk of fiber damages due to the large needles used and the high pressure applied [24]. Therefore, adapting our non-traumatic injection technique for injection of non-toxic vector solution in human nerves via the NHP model is an essential step toward a clinical application of this gene therapy.

The AAV2/9 vector was used to introduce shRNAs into mSC of CMT1A rat sciatic nerves bilaterally in order to decrease PMP22 overexpression. CMT1A rats are transgenic animals that overexpress additional copies of mouse PMP22 in mSC that already express two endogenous copies of rat PMP22 [57]. So we designed two different shRNAs: sh1 significantly decreases both mouse and rat PMP22 mRNA while sh2 targets only mouse PMP22 mRNA. Both shRNAs efficiently and similarly reduced the overall amount of PMP22 in CMT1A rat nerves to that of WT ctr.sh level. As we were unable to distinguish between the mouse and the rat PMP22 proteins, we do not know whether sh1 had a different impact on the mouse or the rat protein expression. However, our data suggest that the reduction of PMP22 protein level is not correlated to the species specificity but to the general amount of active shRNA that is expressed in cells. This indicates that the intensity of the downregulation is positively correlated with the amount of vector injected in the tissue. As PMP22 haploinsufficiency and hence under expression is responsible for peripheral neuropathy with liability to pressure palsy, the control of the silencing intensity represents one of the challenges of this gene therapy. Thus, the definition of the maximal safe dose to be injected is another step toward a clinical application.

We treated young CMT1A animals 6 to 7 days postnatal when peripheral nerve myelination is most active because a large amount of nerve defects and of motor impairments result from the alteration of the initial phase of nerve myelination. Indeed, early nerve defects and impairments already occur in young CMT1A rats [16, 15, 13]. This is consistent with the disease onset occurring in the first decade in 75% of CMT1A patients [40]. Moreover, the treatment of young CMT1A rats (P6 to P18) with soluble Neuregulin-1 was sufficient to halt disease progression at least until 9 weeks of age, while treatment of adult animals has only a limited impact on the disease [16]. We found that PMP22 silencing significantly increased MPZ protein expression in treated CMT1A nerves suggesting that myelin production is increased. Indeed, morphological analysis indicated that significantly more axons were myelinated and the myelin thickness is slightly increased (g-ratio decrease) in treated CMT1A nerves. Moreover, CARS analysis showed that internodes, the myelinated part of the axon between two nodes of Ranvier, were longer in treated CMT1A animals. As the number of myelinated segments and the length of internodes are determined early during myelination [11], taken together this indicates that the deficit of myelination occurring early on in CMT1A rats is corrected by PMP22 silencing following our gene therapy. This is confirmed by the NCV analysis: at one month CMT1A ctr.sh rats already have a reduced NCV compared to WT ctr.sh rats due to the deficit of myelinated segment at early stages of postnatal development. The gene therapy is able to correct this defect as soon as one month, well before defects appear at the motor behaviour level, indicating that this correction occurs directly on the myelination deficit at early stages. While our CARS analysis suggest that PMP22 silencing also prevents late-occurring defects such as focal hypermyelination and segmental demyelination, a benefit of the gene therapy for older diseased animals remains to be shown. Regarding a clinical application, these data suggest that treating CMT1A as early as possible, *e.g* in children, would be more beneficial than in adults. Treating CMT1A children in the long term using a gene therapy approach would constitute a major change as all existing pharmacological strategies target adult patients.

Over the past decade AAV-based therapies have shown numerous successes from proof of concept to clinical trials in genetic diseases [17, 27]. These studies have also identified off-targets transduction and humoral immune response against the vector as the two main serious obstacles that hinder successful AAV-based therapies in patients [39, 61]. We therefore evaluated the biodistribution of AAV2/9 vector throughout the rat body 3 months after injection in sciatic nerves. While 91% of the injected nerves showed vector expression, very few animals had this vector in their liver (12%), heart (9%), kidney (0%), spinal cord (0%), spleen (0%) and blood (14%). Moreover, vector genome copies were detected in dorsal root ganglia L4 and L5 for only few animals (16%) despite the fact that these ganglia are located in the close vicinity of the sciatic nerve. This biodistribution pattern is unusually limited for an AAV2/9-based gene therapy treatment. Indeed, different delivery routes of AAV2/9 in mammals (mouse, rat, dog and NHP), such as intravascular, intracerebroventricular, intrathecal and intraparenchymal routes resulted in a large amount of transduction in the liver and the heart in the majority of injected animals [1, 3, 8, 20, 53, 56]. The intra-nerve injection clearly limits the spread of the vector throughout the body probably through the several layers of cells that compose the nerve surrounding sheath [46]. This restricted biodistribution of the vector will potentially prevent off-target side effects that usually plague gene therapy approaches.

Another probable effect of this limited spread of the vector throughout the body is the low immune response that we observed following the treatment. This response, often directed against the AAV capsid, can block AAV transduction if neutralizing factors are pre-existing. Therefore, the immune response toward the injected therapeutic vector is a serious hurdle for gene therapy treatments when considering vector administration or re-administration *(55, 56)*. Fortunately, among all serotypes, AAV serotype 9 exhibits one of the lowest seroprevalence in humans with less than 50% of the population [5]. In this study, only 20% of the injected animals presented neutralizing factors against AAV2/9 capsid in their blood. Moreover, the titers of these factors were low suggesting they were not abundant. Whether these factors pre-existed the treatment is not known. In any case, we found no correlation between the presence of these factors and a reduced benefit of the therapy suggesting these low-titer neutralizing factors had no impact on the therapy. In addition, these data suggest that a re-injection of the vector in other nerves of treated animals will not generate an acquired immune response to the vector. This opens the possibility for successive treatments of several nerves in CMT1A patients.

A clinical trial of this gene therapy approach will require the measure of the outcome of the treatment. Several clinical scores based on functional tests and assessments such as ONLS [19], CMTNS [41, 49] or CMTPedS for children [7] exist. However, several clinical trials have shown that these scores remain weakly discriminative regarding a slowly progressive disease such as CMT1A [43, 63]. While most of these trials have involved adult patients for whom the disease may progress more slowly than for children, it remains to be seen whether other means are required to measure outcomes. We focused here on skin biomarkers expression described by R. Fledrich and M. Sereda [14, 15]. The expression of several genes in skin biopsies of CMT1A rats were used to identify prognostic and disease severity biomarkers which correlate with clinical impairment. Nine of these genes were then selected as biomarkers of the disease severity in 46 patients [15]. More recently, these biomarkers were further validated in 266 clinically well-characterized genetically proven patients with CMT1A and their use as markers of the disease progression was validated on a 2-3 years interval [14]. We found that, while individually none of these biomarkers was robust enough, taken together they were able to discriminate between sham-treated and PMP22 shRNA-treated animals. Furthermore, we showed that a multi-variated analysis including functional tests (Rotarod, grip strength, NCV and Randall-Selitto) and biomarkers analysis provide a list of 3 biomarkers that are sufficient to measure the outcome of the treatment. A combination of 2 functional tests is also sufficient to cover the full scale of relevant biomarkers. Therefore, we provide here a tool box to reliably measure the outcome of a treatment in a preclinical study in CMT1A rats. The variable number of significant correlations between biomarkers and functional tests also suggests the scalability of the proposed outcome measure. However, this remains to be confirmed in a protocol using different doses of the treatment. In addition, we tested animals 12 months after the treatment. It would be useful to know the robustness of the outcome measure in a less favourable situation such as at one or two months post-treatment. In any case, if one considers that our functional tests in CMT1A rats are similar to clinical scores in patients, our data suggest that combining a clinical score with a transcriptional analysis of the three most relevant skin biomarkers will provide a reliable, robust and probably scalable outcome measure of the gene therapy in CMT1A patients.

## Materials and Methods

### Study design

The goal of this study was to first assess the transduction pattern of AAV vectors serotype 2/9 and 2/rh10 in rodents and NHP after intra nerve injection, and then the efficiency and the safety of a gene therapy approach based on AAV serotype 9 viral vectors expressing shRNA directed against PMP22 mRNA in CMT1A rats. The therapeutic readouts analyzed were: downregulation of PMP22 protein in sciatic nerves using Western blot; upregulation of Myelin Protein Zero level in sciatic nerves using Western blot; nerve fibers morphological evaluation using CARS, immunohistology and thin section electron microscopy thin; nerve electrophysiological analysis; motor and sensory behavioral performances; skin mRNA biomarkers analysis using RT-qPCR; multi-variated analysis including skin biomarkers expression and functional phenotypes data; biodistribution of the vectors in several organs using qPCR; vector neutralizing factors tittering in the blood. Experimental groups were sized according to the literature to allow for statistical analysis. No outliers were excluded from the study. Behavioral data originating from animals that died or had physical disabilities unrelated to CMT1A disease during the study were not used. Sample collection, tissue processing and treatment are included and described in the Results and Materials and Methods. Rats were randomly assigned to the different experimental groups after genotyping. Scientists who performed the experiments and analysis were blinded to the group’s identity. Data were analyzed by those carrying out the experiments and verified by the supervisors.

### Cloning and vector production

Cloning of the transgenes in pAAV and AAV vector productions were provided by the CPV Vector Core of INSERM UMR 1089, Université de Nantes (France). Briefly, ssAAV2/9-CAG-eGFP and ssAAV2/rh10-CAG-eGFP vectors were obtained from pAAV-CAG-eGFP plasmid containing AAV2 inverted terminal sequences, CAG promoter and the enhanced green fluorescence protein (eGFP). These vectors were used to determine the transduction pattern following a direct intra-nerve injection. ShRNA sh1 (5’-CGCGGTGCTAGTGTTGCTCTT-3’) recognizes both Rattus Norvegicus and Mus Musculus PMP22 mRNAs while shRNA sh2 (5’-CACTGACTACTCCTATGGCTT -3’) only recognizes Mus Musculus PMP22 mRNA. Both shRNAs were cloned under the control of U6 promoter in a pAAV plasmid expressing eGFP protein under a CAG promoter (pAAV-sh1 and pAAV-sh2 respectively). These 2 plasmids were used to generate AAV2/9-sh1 and AAV2/9-sh2 vectors. These 2 vectors were used to evaluate their efficiency in the CMT1A rat model. A control vector AAV2/9-U6-ctrl.sh-CAG-eGFP expressing a shRNA with no target in mammals served as a control (named AAV2/9-ctr.sh).

Vector production was performed following the CPV facility protocol as previously described [2]. Briefly, recombinant AAVs were manufactured by co-transfection of 293-HEK cells and purified by cesium chloride density gradients followed by extensive dialysis against phosphate-buffered saline (PBS). Vector titer were determined using a vector-specific qPCR assay.

### Animals included in this study

All animal experiments were approved by the local ethic committee and the ministère de la recherche et de l’enseignement supérieur (authorization 2017032115087316 and 2016091313354892 for rodents and 2015061911295753v2 for NHP). All the procedures were performed in accordance with the French regulation for the animal procedure (French decree 2013-118) and with specific European Union guidelines for the protection of animal welfare (Directive 2010/63/EU).

#### Mice

C57BL/6 mice were purchased from Janvier Labs (France). These mice were used to evaluate the transduction pattern of AAV2/9 and AAV2/rh10 after intra-nerve injection. A cohort of 3 adult mice (2-3 months old) and 3 pups (P2-P3) were injected with each AAV vectors as described in the section “vector delivery” below.

#### Rats

A breeding colony was established from a gift of CMT1A rats from Max Planck Institute of Experimental Medicine, Goettingen, Germany. Littermates and CMT1A rats were identified using PCR on DNA isolated from the tail as described previously [57]. For the transduction pattern, a cohort of 3 adult rats (2-3 months old) and 3 pups (P6-P7) were injected with each AAV vectors as described in the section “vector delivery” below.

For the gene therapy assay, wild-type littermate (WT) and CMT1A male and female rats were randomly divided into 4 groups (WT ctr.sh, CMT1A ctr.sh, CMT1A sh1, CMT1A sh2) of 16 rats each. At 3 months of age half of the rats in each group (8 per group) were sacrificed for biochemical and biodistribution studies. All the others were kept until 12 months of age to study the efficiency of the gene therapy on the sensory-motor behavior and NCV at different time points post injection (1, 2, 3, 6, 9 and 12 months). Finally, these animals were sacrificed for histological and biochemical studies.

#### Nonhuman Primates

Two juvenile cynomolgus macaques (*Macaca fascicularis*, two females, 3.7 years old/ 4.3 kg and 2.3 years old/2.9 kg) were included in the study. Animals were part of the the MIRCen colony (CEA Fontenay-aux-roses, France). Progenitors were imported from a licensed primate breeding centers on Mauritius and Philippines. The experiments were performed in an authorized user facility (Ministère de l’Agriculture, number 92–032-02). Monkeys remained under veterinary care during the full study. Animals were tested negative for anti-AAV2/9 or anti-AAV2/rh10 antibodies before the treatment.

### Vector delivery

#### Mice and rats

The intra-nerve injection of AAV vectors into the sciatic nerve was performed as previously described [18]. Briefly, under anesthesia with isoflurane, the skin on the thigh was disinfected with betadine solution and ethanol 70% and cut above the sciatic nerve location. *Biceps femoris* and *gluteus superficialis* muscles were carefully separated to expose the small cavity containing the sciatic nerve which was lifted with a spatula. Next, the viral solution was injected using a fine glass needle (borosilicate glass capillary GC 100-10, Harvard Apparatus, France) manipulated with a micromanipulator (World Precision Instruments, France). The injection was performed using a pneumatic picopump (PV820, World Precision Instruments, France) controlled by a pulse generator (GW GIG8215A, INSTEK, France). At the end of injection, the sciatic nerve was placed back inside its cavity, muscles were replaced and the wound was closed with staples (12-020–00, Fine Science Tools, France). Animals were then treated with buprenorphine (100 μg/kg) for two days.

AAV vector solutions were prepared by diluting vectors at the right titer with sterile phosphate-buffered saline and 0.01% fast green. Pups and adult mice were unilaterally injected with 2 µl containing 1 × 10^10^ vg and with 8 µl containing 5 × 10^10^ vg respectively. Pups and adult rats were injected unilaterally or bilaterally with 1 × 10^11^ vg/nerve in 8 µl and 1.8 × 10^11^ vg/nerve in 30 µl respectively.

#### Nonhuman Primates

Animals were sedated through an intramuscular treatment of ketamine hydrochloride (Imalgene, 10 mg/kg) and xylazine (2% Rompun, 0.5 mg/kg). Anaesthesia was then maintained with propofol (Propovet, 10 mg/kg/h, intravenous infusion in the external saphenous vein).

After anaesthesia, animals were placed in prone position and a small incision on the skin was performed above the popliteal fossea. Animals were injected in the sciatic nerves 1 cm above the bifurcation between common fibular and tibial nerves. Both animals received a single and unilateral injection in the left sciatic nerve. A 22G needle was manually inserted under the epineurium to guide a 150µm-wide silica capillary 3 mm deep into the nerve. The capillary was connected to a 1ml Hamilton syringe mounted on an infusion pump (Harvard Apparatus, France) to allow infusion of the vector at 13.9μl / min. NHP were injected with 5 × 10^12^ vg/nerve in 416µl. Vectors were diluted in Fast green (0.005% final concentration). During the procedure, animals were placed on warming blankets and physiological parameters were monitored. After full recovery from anesthesia, they were replaced in their home cages and clinical observations were daily performed by trained technicians for any sign of discomfort or distress during seven days.

### Tissue collection and processing

#### Mice and Rats

Animals were euthanized one month (transduction pattern study) or 3 months and 12 months (gene therapy assay) post-injection using Pentobarbital (54.7 mg/mL, 100 mg/kg, CEVA Santé Animale, France). They were transcardially perfused with phosphate-buffered saline (PBS) and sciatic nerves were freshly and quickly dissected. Nerves were then fixed for 1h in 4% paraformaldehyde (PFA)/PBS solution at room temperature or 24h in 4% PFA/ 2.5% glutaraldehyde/PHEM (PIPES, HEPES free acid, EGTA, MgCl2) for electron microscopy analysis. For biochemical studies, tissues were directly frozen in liquid nitrogen and stored at -80°C.

#### Nonhuman Primates

Euthanasia was achieved through an overdose of Pentobarbital Sodium (180 mg/kg, intravenous injection), four weeks after injection, under sedation by intramuscular injection of ketamine hydrochloride (Imalgene, 10 mg/kg) and xylazine (2% Rompun, 0.5 mg/kg). Animals were then intracardially perfused with 1 L of PBS then 2.5 L of ice cold 4 % PFA/PBS. For each NHP, the injected sciatic nerve was examined and collected. Contro-lateral non injected sciatic nerves served as a control. After one night of post-fixation in 4% PFA/PBS, they were processed for histological studies as described in the histological study section.

### Histological study

Mouse anti-Myelin Basic Protein (MBP-SMI-99, Millipore, reference NE1019, 1/1000 dilution), rat anti-Myelin Basic Protein (BIO-RAD, reference MCA409S, 1/1000 dilution), rabbit anti-β-Tubulin III (Tuj1, Sigma reference T2200, 1/1000 dilution), donkey anti-mouse Alexa Fluor 594 (Thermofisher, reference A-21203, 1/1000), donkey anti-rat Alexa Fluor 594 (Thermofisher, reference A-21209, 1/1000) and donkey anti-rabbit Alexa Fluor 647 (Thermofisher, reference A-31573, 1/1000 dilution) were used for both immunohistochemistry on frozen sections and on teased fibers.

#### Immunohistochemistry on frozen sections

Following fixation, samples were incubated 24-48h in two successive baths of 6% and 30% sucrose and then embedded in Optimal Cutting Temperature (OCT, NEG-50, MM France) and stored at -80°C. Coronal sections (10µm of thickness) were cut using cryostat apparatus (LEICA CM3050). Cryosections were blocked with 5% Normal Goat Serum (NGS) and 0.1% triton X-100 in PBS, incubated overnight at 4°C with primary antibodies diluted in NGS/Triton/PBS, washed with PBS and then incubated 1h at RT with secondary antibodies diluted in NGS/Triton/PBS. After several PBS washes, cryosections were mounted in Dako fluorescent mounting medium (S3023). AxioScan slide scanner (Zeiss, France) was used to obtain images. The percentage of transduced myelinating Schwann cells over all myelinating Schwann cells in the full section was calculated using Zen software (Zeiss, France).

#### Immunohistochemistry on teased fibers

After sciatic nerve fixation, fibers were gently teased, dried on glass slides, and stored at -20°C. Teased fibers were permeabilized by immersion in –20°C acetone for 10 minutes, blocked at room temperature for 1 hour with NGS/Triton/PBS, then incubated overnight at 4°C with primary antibodies diluted in NGS/Triton/PBS. Slides were then washed several times with PBS and incubated with secondary antibodies diluted in NGS/Triton/PBS. Slides were mounted on coverslips with Dako fluorescent mounting medium (S3023) and examined using an Apotome fluorescence microscope (Zeiss, France). For each sciatic nerve, 10 fields were captured and the percentage of myelinating Schwann cells, non myelinating Schwann cells and axons transduced by AAV vectors among all transduced cells were calculated using Zen software (Zeiss, France).

#### Electron microscopy of sciatic nerve

Fixed nerves were then processed by the Electron Microscopy platform at the Institute of Neurosciences in Montpellier (INM, MRI-COMET). Samples were post-fixed with osmium 0.5% and K4FeCN6 0.8% 24h at RT in the dark, went through dehydration in successive ethanol baths and embedding in Epoxy resin (EMbed-812 Embedding Kit, Electron Microscopy Sciences, France) using a Leica AMW machine. They were finally incubated 36 h in an oven at 60°C. Nerves were cut with Leica Reichert Ultracut S ultramicrotome into semi-thin sections of 700nm-1µm. Sections were stained with Toluidine Blue and imaged using an AxioScan slide scanner (Zeiss, France). The total number of myelinated axons, the mean axon diameter (200 axons per nerve randomly distributed throughout the entire section) and the g-ratio (axon diameter over the full fiber diameter) were measured using ImageJ software and GRatio plugin.

### Cell culture and transfection

The efficiency of PMP22 sh1 and sh2 was examined *in vitro* using Schwann cell lines from mouse (MSC 80) (JM Boutry et al, J Neuro Research, 1992) and rat (RT4-D6P2T). 500 000 cells per well were seeded in a 6-well-plate in Dulbecco’s Modified Eagle Medium (DMEM) (Gibco/Thermo Fisher, France) with 10% Fetal Bovine Serum (FBS) (Gibco/Thermo Fisher, France) and 1% Penicillin Streptomycin (Gibco/Thermo Fisher, France). Twenty-four hours after seeding, cells were transfected using jetPRIME reagent (Polyplus-transfection S.A, France) according to manufacturer protocol with pAAV-ctr.sh or sh1 or sh2. Two days after transfection, each well was washed 3 times by PBS and proteins were extracted using RIPA lysis buffer completed by protease inhibitors (Fisher Scientific, France) for 1h at 4°C. Three independent experiments were performed with each cell line.

### Western blot

Frozen nerves were crushed with a pestle and mortar cooled on dry ice, solubilized in RIPA lysis buffer completed with protease inhibitors (Fisher Scientific, France) and homogenized on a rotating wheel at 4 °C for 3 hours. They were then sonicated 3 times during 10 seconds on ice (Microson ultrasonic cell disruptorXL, Microsonic) and centrifuged for 30 minutes at 10,000rpm at 4°C. Proteins concentrations were quantified using the Bicinchoninic acid (BCA) protein assay kit (Thermo Scientific, France). Ten micrograms of proteins was loaded on a 4-20% precast polyacrylamide gels (Mini Protean gels, Bio Rad, France). Then, proteins were transferred to nitrocellulose membranes (Bio Rad Trans blot transfer pack) through semi-dry transfer process (Bio Rad Bio Rad Trans-Blot Turbo system). Membranes were first incubated with the REVERT total protein stain solution (LI-COR Biosciences, France) to record the overall amount of protein per lane. Then, membranes were blocked for 1 hour at room temperature using LI-COR blocking buffer (Odyssey Blocking buffer, LI-COR Biosciences, France). They were then incubated with the following primary antibodies overnight, at 4°C in the same blocking buffer: rabbit anti-PMP22 (Sigma-Aldrich, SAB4502217, 1/750 dilution), mouse anti-β-Actin Clone AC-15 (Sigma-Aldrich, A1978, 1/10 000 dilution) or goat anti MPZ (Thermo Fisher Scientific, PA5-18773, 1/1000 dilution). Following 3 washes of 10 min with TBS-0.1% Tween (TBST), secondary antibodies were used at a 1/15 000 dilution in Li-COR blocking buffer: IRDye 800CW donkey anti-rabbit (LI-COR Biosciences, 925-32213), IRDye 680RD donkey anti-mouse (Li-COR Biosciences, 925-68072) or IRDye 800CW donkey anti-goat (LI-COR Biosciences, 925-32214). After 3 washes in TBST, results and quantifications were obtained by the Odyssey CLX LI-COR Imaging System and its “Image Studio” software.

### CARS imaging

All CARS images were obtained as described previously [21] with a two-photon microscope LSM 7 MP coupled to an OPO (Zeiss, France) and complemented by a delay line [42]. A ×20 water immersion lens (W Plan Apochromat DIC VIS-IR) was used for image acquisition. Four consecutive fields (250mm square, 200µm deep) were captured on sciatic nerves and an image of 1cm long / 20µm deep was then reconstructed using Zen software (Zeiss, France). The number of nodes of Ranvier was determined for each reconstructed image. The results were expressed as a density of nodes of Ranvier over the number of fibers per field.

### Behavioral analysis

#### Rotarod

The rotarod test was performed using the “Rota Rod” machine specific for rats (Bioseb, France). Following one day of training, the latency to fall from the rotating bar was recorded the next day with an acceleration from 4 to 40 rpm over a period of 5 minutes. Each animal underwent three trials. Data were averaged for each rat and for each group.

#### Griptest

The grip test was performed on the rear paws using a grid connected to an electronic device recording the force in Newtons (Bioseb, France). The muscular strength was measured by recording the maximum amount of force maintained by the animal after it gripped the grid with its rear paws and while it was pulled down by its tail. Each animal underwent three trials. Data were averaged for each rat and for each group.

#### Randall Selitto paw-pressure vocalization Test

Mechanical pain sensitivity was measured as the threshold to a noxious mechanical stimulus as previously described [48]. Briefly, for 2 weeks before the experiments, animals were daily placed in the experimental room and were left for 1 h to become accustomed to the environment. Then, they were gently handled for 5 min and exposed to the nociceptive apparatus without stimulation. To evaluate the nociceptive threshold, a constantly increasing pressure was applied to the rat hind paw until vocalization occurs. A Basile analgesia meter (stylus tip diameter, 1 mm; Bioseb, France) was used. A 600-g cut-off value was determined to prevent tissue damage. Results are expressed as the mean of each hindpaw per rat.

### Electrophysiology analysis

Nerve conduction velocity (NCV) was measured on both sciatic nerves of anaesthetized rats (isoflurane, AST-00 manually operated workstation, Anesteo, France) placed on a heating plate at 37° C. Proximal and distal stimulations were performed using a pair of 12 mm-steel needle electrodes with 2 mm pin plugs (AD Instruments, MLA1304, Oxford, UK) and were recorded from the intrinsic foot muscles using 12 mm-steel electrodes with 1.5 mm safety socket plugs (AD Instruments, MLA1303, Oxford, UK) placed on the rat’s rear paw’s plantar muscle and on the middle toe’s muscle. A biphasic stimulation lasting 0.2 ms was applied using a PowerLab 26T generator (AD Instruments, Oxford, UK) connected to LabChart software (AD Instruments, Oxford, UK). Stimulations were delivered with an increasing current intensity until supramaximal stimulation. NCV was calculated as the ratio of the distance between the proximal and distal sites of stimulation in meter on the action potential latency in second using LabChart software.

### Biomarkers studies

Glaber skin of front paws were collected on animals sacrificed 12 months post-injection. Samples were snap-frozen in liquid nitrogen and stored at - 80 °C for biomarkers analysis as described previously [14]. Briefly, total RNA was extracted using RNeasy Mini Kit (Qiagen, France) according to manufacturer’s recommendations for non-fatty tissue and precipitated in ice-cold ethanol. The Agilent integrity check was used to verify RNA quality. Samples with an integrity number higher than 7 were used for cDNA synthesis using the Superscript III RT kit (Invitrogen, Germany). Real-time semiquantitative PCR with TaqMan and SYBRGreen were performed in the LightCycler 480 Systems (384-well format, Roche Applied Science, Germany) with a reaction mix prepared to the final volume of 10 µL. All reactions were run in four replicates. The threshold cycles for each gene of interest were normalized against two stable housekeeping genes (*ActB* and *Rplp0*). The PCA and correlation matrix were performed using R-package ade4.

### AAV2/9 biodistribution

Samples were collected from rats 3 months after injection. We collected sciatic nerves at the injection site, the lumbar dorsal root ganglia 4 and 5 (DRG L4 and L5), the lumbar spinal cord, the heart, the liver, the spleen, the kidney, the brainstem and the blood. Whole blood was collected in tubes containing EDTA. All samples were collected in DNA-free, RNAse/DNAse-free and PCR inhibitor-free certified microtubes. Tissue samples were obtained immediately after sacrifice in conditions that minimize cross-contamination and avoid qPCR inhibition, as described previously [32]. Samples were snap-frozen in liquid nitrogen and stored at -80°C. Extraction of genomic DNA (gDNA) from 200µL of whole blood or from tissues using the Gentra Puregene kit and Tissue Lyser II (Qiagen, France) was performed accordingly to manufacturer recommendations.

qPCR analyses were conducted on a StepOne Plus apparatus (Applied Biosystems®, Thermo Fisher Scientific, France) using 50ng of gDNA in duplicates. All reactions were performed in a final volume of 20µL containing template DNA, Premix Ex Taq (Ozyme, France), 0.4µL of ROX reference Dye (Ozyme, France), 0.2µmol/L of each primer and 0.1µmol/L of Taqman® probe (Ozyme, France). Vector genome number was determined using the following primer/probe combination, designed to amplify a specific region of the *GFP* transgene:

Forward: 5’-ACTACAACAGCCACAACGTCTATATCA-3’,

Reverse: 5’-GGCGGATCTTGAAGTTCACC-3’,

Probe: 5’-FAM-CCGACAAGCAGAAGAACGGCATCA-TAMRA-3’

Endogenous gDNA copy numbers were determined using the following primers/probe combination, designed to amplify the rat *Hprt1* gene:

Forward: 5’-GCGAAAGTGGAAAAGCCAAGT -3’,

Reverse: 5’-GCCACATCAACAGGACTCTTGTAG-3’,

Probe: JOE-CAAAGCCTAAAAGACAGCGGCAAGTTGAAT-TAMRA-3’.

For each sample, threshold cycle (Ct) values were compared with those obtained with different dilutions of linearized standard plasmids (containing either the *GFP* expression cassette or the rat *Hprt1* gene). The absence of qPCR inhibition in the presence of gDNA was checked by analyzing 50ng of gDNA extracted from tissues samples or blood from a control animal. Results were expressed in vector genome number per diploid genome (vg/dg). The limit of quantification (LOQ) was determined as 0.002 vg/dg.

### AAV2/9 Neutralizing factors

The detection of AAV2/9 neutralizing factors was performed by the Gene Therapy Immunology Core (GTI) at INSERM UMR 1089 laboratory (Nantes, France). Briefly, the neutralization test consisted in an *in vitro* cell transduction inhibition assay using a standard recombinant AAV2/9 expressing the reporter gene system LacZ. Serum samples were collected from rats 3 months post injections. Each serum was tested using a range of dilutions 1/50, 1/500, 1/5000, 1/50 000 and 1/500 000. Gene expression was measured in cell lysates using a chemoluminescent substrate of galactosidase (GalactoStar kit, Invitrogen). Neutralizing factor titer corresponds to the last dilution leading to the inhibition of more than 50% of the standard AAV2/9 transduction (100% corresponds to the maximum transduction obtained with the standard AAV2/9 alone). Five rats injected with AAV2/9-ctr.sh and fives CMT1A rats injected with AAV2/9-ctr.sh were randomly analyzed for the detection of AAV2/9 neutralizing factors.

### Statistical analysis

Data were analyzed with Graphpad Prism version 7 (Graphpad Software) and expressed as the mean ± standard error of the mean (SEM) or ± standard deviation (SD) as indicated in the Figures legends. Statistical differences between mean values were tested using one-way or two-way ANOVA analysis followed by Tukey’s or Dunnett’s multiple comparison test as indicated in the Figures legends. Differences between values were considered significant with: \**P* < 0.05, ** *P* < 0.01, *** *P* < 0.001, **** *P* < 0.0001.

## Acknowledgments

We thank the SMARTY platform (RAM - Animal facility network of Montpellier) for expert care of animals and help for behavioral tests (Anne-Laure Bonnefont), the Electron Microscopy platform (Chantal Cazevieille), Montpellier Ressources Imaging (Hassan Boukhaddaoui), CEA MIRCen Institut (CEA Fontenay aux Roses, France), CPV vector core and Gene Therapy Immunology core from the INSERM UMR 1089, University of Nantes (https://umr1089.univ-nantes.fr/facilities-cores/). We also are grateful to Klaus-Armin Nave and Mickael Sereda for their gift of CMT1A rat.

## Author contributions

Conceptualization: BG, NT; Formal analysis: BG, HH, RF; Funding acquisition: PA, NT; Investigation: BG, HH, SS, JB, MD, SA, GC, CMF, AJ, CR, MZ, VFLR, CLG, RF; Methodology: HH. Project administration: BG, NT; Resources: BG, CMF, VFLR, CLG, AJ, CR, PA, RF; Supervision: NT; Validation: BG, HH, NT; Visualization: BG, HH, NT; Writing – original draft: BG, HH, NT; Writing – review & editing: all co-authors.

## Data and materials availability

All data and materials are available upon request.

## Electronic Supplementary Material

**ESM 1** Representative performance of a CMT1A ctr.sh rat on a narrow beam 12 months after injection

**ESM 2** Representative performance of a CMT1A sh1 rat on a narrow beam 12 months after injection

**ESM 3** Transcriptomic analysis of nine biomarkers in the front paw skin

