## Supplementary Material 3 for "Long term AAV2/9-mediated silencing of PMP22 prevents CMT1A disease in rats and validates skin biomarkers as treatment outcome measure"

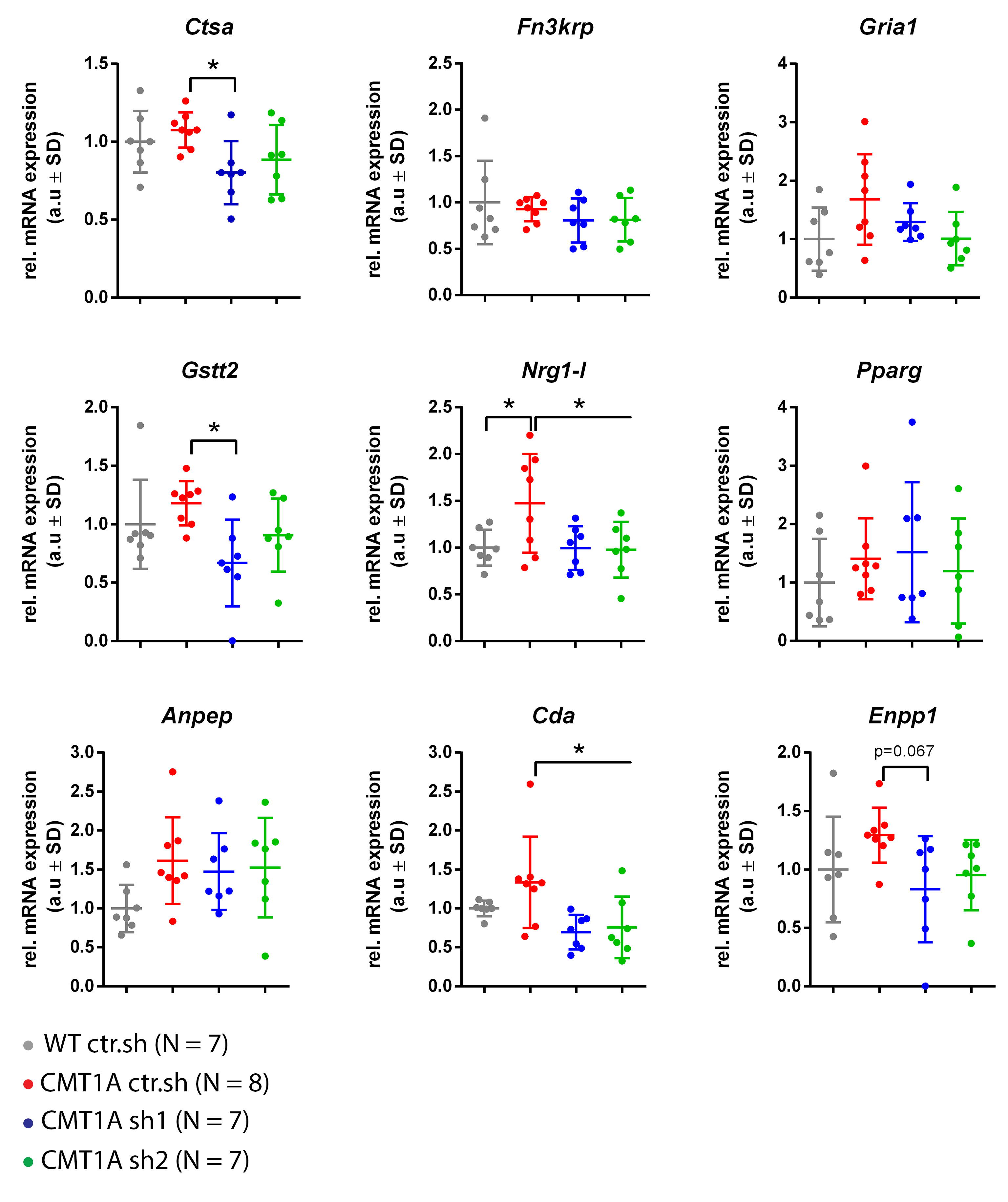

**ESM 3** Transcriptomic analysis of nine biomarkers in the front paw **skin. The** mRNA expressions of nine biomarkers (label on the top of each graph) were quantified using RT-qPCR in the skin of front paws and normalized against the mRNA expression levels of two stable housekeeping genes. Relative (rel.) mRNA expressions are shown in arbitrary unit (a.u). Error bars show SD. Statistical tests show one-way ANOVA followed by Dunnett’s post hoc test. * *P* < 0.05. When statistical test results are not shown, this indicates non-significant results.
